# Ancestral polyploidy and non-linear dosage shifts define the photosynthetic transition in the Heliantheae alliance

**DOI:** 10.64898/2026.11.05.764543

**Authors:** Venkata Suresh Bonthala, Konoutan M. Kafoutchoni, Stephanie Krey, Urte Schlüter, Andreas P.M. Weber, Dimitri Tolleter, Benjamin Stich

**Affiliations:** Julius Kühn-Institut (JKI), Institute for Breeding Research on Agricultural Crops, Rudolf-Schick-Platz 3a, OT Groß Lüsewitz, Sanitz, 18190, Germany; Institute of Quantitative Genetics and Genomics of Plants, Faculty of Mathematics and Natural Sciences, Heinrich Heine University, Düsseldorf, Germany; Institute of Plant Biochemistry, Faculty of Mathematics and Natural Sciences, Heinrich Heine University, Düsseldorf, Germany; Biosphere Science and Engineering Division, Carnegie Science, Stanford, CA 94305, USA

**Keywords:** Whole-genome duplications, gene copy number variations, CO_2_ compensation points, carbon concentrating mechanisms, Heliantheae alliance

## Abstract

Whole-genome duplications (WGDs) have played a central role in plant evolutionary innovation, yet their genome-wide influence on complex physiological traits such as carbon-concentrating mechanisms (CCMs) remains poorly resolved. Here, we present high-quality genome assemblies and annotations for 35 species within the Heliantheae alliance (Asteraceae), spanning the complex photosynthetic spectrum from C_3_ to C_4_. Ks-based analyses combined with gene tree-species tree reconciliation identified a shared paleopolyploid event (WGD-2) ∼30 million years ago. Using phylogenetically informed association mapping with these species, we detected 451 functional MapMan BINs with copy number variation significantly associated with interspecific variation in the CO_2_ compensation point. Notably, 303 of these BINs contain ≥10% WGD-2-derived genes, indicating a strong enrichment of duplicated loci within photosynthetically relevant functional modules. Effect-size analysis revealed that the C_3_-to-C_4_ transition is governed by threshold-dependent genomic architecture rather than gradual additive change. Substantial expansions in key transcription factor modules—including HD-ZIP, HSF, LOB, and TCP—were associated with strong non-linear physiological shifts, consistent with genomic tipping points. By integrating comparative genomics, physiological trait variation, and evolutionary analysis, this study establishes a mechanistic link between ancient polyploidy and contemporary photosynthetic diversification. These findings provide a framework for understanding the genetic basis of carbon concentrating mechanism evolution and inform future optimization of photosynthetic efficiency in Asteraceae crops such as sunflower.

## Introduction

Photosynthesis serves as the primary metabolic conduit for the synthesis of nearly all biomass on earth (Evans, 2013). At the molecular level, this process is driven by highly-conserved multi-protein complexes that execute photon capture, electron transport, and enzymatic carbon assimilation through the Calvin-Benson cycle (Eberhard et al., 2008). This complex, multi-stage process has been shaped by numerous independent evolutionary trajectories (Sage, 2004). Over time, these trajectories have refined carbon-fixation strategies across diverse plant lineages. However, the ancestral C_3_ pathway remains the most prevalent mode of carbon assimilation (Ehleringer et al., 1997). Despite its dominance so far, it is fundamentally constrained by the oxygenase activity of ribulose-1,5-bisphosphate carboxylase/oxygenase (Rubisco) under current atmospheric CO_2_ concentrations (Bowes et al., 1971; Walker et al., 2016). Specifically, Rubisco frequently mistakes O_2_ for CO_2_, producing the toxic metabolite 2-phosphoglycolate rather than sugar-building intermediates. This constraint gives rise to the energy-intensive photorespiratory cycle, which salvages 2-phosphoglycolate (Ogren, 1984; Sharkey, 1988). Although this pathway is metabolically necessary, it significantly reduces photosynthetic efficiency and biomass accumulation.

To circumvent these constraints, various plant lineages have independently evolved carbon-concentration mechanisms (CCMs). Most notably, C_4_ photosynthesis has arisen in more than 60 distinct instances of convergent evolution (Sage, 2004; Sage et al., 2011, 2012). This sophisticated adaptation relies on the spatial compartmentalization of carbon fixation. By elevating CO_2_ levels in spatial proximity of Rubisco, it suppresses oxygenase activity and thereby enhances carboxylation efficiency. Achieving these specialized CCMs requires coordinated anatomical and biochemical modifications. These include the development of Kranz anatomy and the recruitment of key enzymes such as phosphoenolpyruvate carboxylase (PEPC) and pyruvate, orthophosphate dikinase (PPDK) (Brown, 1975; Hatch, 1987; Lundgren et al., 2014; Sage et al., 2014).

Gene duplications, whether single-gene or whole-genome events, are recognized as critical genomic prerequisites for the transition of C_3_ to C_4_ photosynthesis (Monson, 2003). This pattern has been documented primarily for core enzymes in Flaveria (Ludwig, 2011; Schulze et al., 2013), Alloteropsis (Bianconi et al., 2018), and Cleome (Huang et al., 2021; Hoang et al., 2023). For example, the photorespiratory GLDP gene underwent duplication in the ancestral C_3_ Flaveria species (Schulze et al., 2013). One copy subsequently evolved predominant bundle-sheath-specific expression. This regulatory divergence facilitated the emergence of a photorespiratory CO_2_ pump in C_3_–C_4_ intermediate Flaveria. Similarly, orthologs of several essential C_4_ cycle enzymes—including PEPC, PPDK, NADP-ME, NADP-MDH, and CA— were duplicated in C_3_ lineages in Flaveria. These duplications created a genomic substrate in which one paralog was free to undergo neofunctionalization. In parallel, dosage-driven upregulation increased enzymatic capacity. Together, these changes supported the evolution of the specialized NADP-ME CCM (Monson, 2003; Ludwig, 2011; Bianconi et al., 2018). Collectively, the above-mentioned studies demonstrate that gene duplication promotes both subfunctionalization and neofunctionalization (Blanc & Wolfe, 2004; Cusack & Wolfe, 2007; Panchy et al., 2016). Together, these mechanisms enable cell-type-specific expression and dynamic regulatory changes that drive the specialization of photosynthetic functions. However, these few well-characterized biochemical components explain only a fraction of the C_4_ phenotype. The broader suite of regulatory and structural genes required to orchestrate a fully integrated CCM remains largely unidentified.

WGDs have been mechanistically linked to a wide range of key adaptations. These include morphological shifts, such as fruit development in tomato (Tomato Genome Consortium, 2012) and fiber quality in cotton (Li et al., 2015a). They have also been associated with complex physiological traits, including nitrogen fixation (Young et al., 2011) and stress tolerance (Huang et al., 2020). Additional examples are reviewed by Qiao et al (2022). Despite this extensive catalog of phenotypic impacts, the specific functional consequences of WGDs for photosynthetic performance remain underexplored.

The Asteraceae family offers a unique system for investigating the genomic basis of photosynthetic evolution as related genera exist next to the crop sunflower—such as *Flaveria*, *Parthenium*, and *Pectis*—that collectively encompass the full range of photosynthetic types: C₃, C₃–C₄ intermediates, C₄-like, and C₄ (Moore et al., 1987; McKown & Dengler, 2007; Hansen et al., 2016; Siniscalchi et al., 2021; Adachi et al., 2023). This natural diversity across species within a narrow phylogenetic context creates an opportunity to test how ancient genome duplications (WGDs) may have contributed to the emergence of CCMs. In particular, it allows us to examine the so far disregarded question whether interspecies variation in gene dosage played a role in this process and this also for genes outside of the core photosynthetic machinery.

In this study, we (1) established genomic resources for evolutionary analysis of photosynthesis in Asteraceae, (2) identified 451 MapMan BIN functional categories by interspecies genome-wide association analysis, and (3) validated the associations in summary using prediction models. Our results indicate that ancient WGD events shaped specific gene copy number architectures. These architectures closely align with the emergence of photosynthetic phenotypes. Together, these patterns point to a genomic mechanism that likely contributed to CCM adaptation. By mapping dosage-sensitive shifts, we clarify how gene copy number variation relates to photosynthetic evolution. These findings highlight potential strategies for enhancing photosynthetic efficiency in sunflower, *Guizotia abyssinica* (Padhi & Panda, 2026) and its related species.

## Results

### Genome assemblies and annotations

In this study, we analysed a total of 38 species across 11 genera within the Asteraceae family. We generated 35 new high-quality genome assemblies specifically for this work (Supplementary Table S1). These represent eight genera from the Heliantheae alliance and two outgroup genera: Artemisia (Anthemideae tribe) and Pallenis (Inuleae tribe). To broaden the genotypic space, this dataset was supplemented with three additional high-quality genomes: *Flaveria bidentis* from our previous study (Sun et al., 2025), and the publicly available assemblies of *Flaveria brownii* (Taniguchi et al., 2021) and *Helianthus annuus* (Badouin et al., 2017).

We estimated genome sizes for the 35 newly sequenced species using a k-mer-based approach. Overall, these k-mer-based estimates were generally consistent with values reported in the literature obtained via flow cytometry and Feulgen densitometry (Taniguchi et al., 2021; https://www.asteraceaegenomesize.com), though some deviations were observed. For example, we estimated the genome size for *Coreopsis lanceolata* as 1.92 Gb, while previous estimates based on flow cytometry (https://www.asteraceaegenomesize.com) recorded a larger size of 3.359 Gb (Supplementary Table S2).

Reference-quality assemblies were generated for nine of the 35 newly sequenced species using the DeNovoMAGIC pipeline (NRGene, Nes Ziona, Israel), yielding an average N50 of over 14 Mb and capturing over 95% of BUSCO genes (Seppey et al., 2019). The remaining 26 species were assembled at draft quality using Supernova v2.1.1 (Weisenfeld et al., 2017) and Flye v2.9 (Kolmogorov et al., 2019), achieving an average N50 of 528 kb and 91.8% BUSCO (Seppey et al., 2019) completeness (Supplementary Figures S1–S2 and Table S3).

Repetitive elements were annotated in all 35 newly assembled genomes and the genome assembly of *Flaveria bidentis* using EDTA v2.0.0 (Ou et al., 2019), with repeat content ranging from 48.58% (*F. robusta*) to 80.46% (*A. vulgaris*; Supplementary Figure S1). Long-terminal repeat (LTR) retrotransposons (Copia, Gypsy, and other LTRs) made up on average across all species the largest fraction (average 46.78%), followed by Helitrons (9.64%) and Mutators (4.33%) (Supplementary Figure S3 and Table S4). Gene annotation was performed in all 35 newly assembled genomes and the genome assembly of *F. brownii* using the deep learning-based tool Helixer v0.3 (Stiehler et al., 2021), yielding 27,179 to 80,151 predicted gene models per species, with an average length of 3.2 kb (Supplementary Figure S4 and Table S3). Functional annotation assigned PFAM domains, Gene Ontology (GO) terms using InterProScan (Jones et al., 2014) and MapMan BIN classifications (Schwacke et al., 2019) to an average of 76%, 50.7% and 93% of genes, respectively, demonstrating consistent annotation depth across all species. An average of 75.68% of the genes showed detectable gene expression (TPM >=0.5) in at least one tissue or pooled tissues (TPM > 0) using direct RNA sequencing, thereby validating the structural accuracy and coding potential of the Helixer-derived gene models (Supplementary Figure S4 and Table S3). Consistent with broader patterns in plant genome evolution (Badouin et al., 2017), genome size variation across Heliantheae alliance was primarily driven by repeat content (R² = 0.995) rather than gene number (R² = 0.544; Supplementary Figure S5).

### Evolutionary relationships among species and dating speciation

We identified orthologous gene families across the genome sequences of all available 38 species using OrthoFinder v2.5.5 (Emms & Kelly, 2019), assigning 96.36% of all genes (1,339,739) to 55,238 phylogenetic hierarchical orthogroups (HOGs). Gene assignment rates varied among species, ranging from 86.98% in *H. annuus to* 99.11% in *F. australasica*, with an average of 96.73%. The number of HOGs per species varied from 19,514 in *C. maritima* to 26,411 in *A. vulgaris* (Supplementary Figure S6 and Table S5). On average, 25.68% of HOGs per species were shared by all 38 species, representing highly conserved gene families. Over 1% of HOGs per species were specific to a single species (species-specific HOGs), while 73.25% were shared among a subset of species (Supplementary Figure S6 and Table S5). These findings indicate both deep conservation and lineage-specific evolution of gene families. Notably, *A. vulgaris*, one of two above mentioned outgroup species, exhibited the highest proportion of species-specific HOGs (7.8%), followed by *C. diversifolius* (5.5%), potentially reflecting recent adaptive evolution or genome duplication events. Pairwise orthologous gene sharing averaged 84.9% between species (Supplementary Figure S7).

To infer species relationships on a genome-wide level, we reconstructed phylogenetic trees based on the maximum likelihood (ML) method with RAxML-ng v1.2.2 (Kozlov et al., 2019) and the multi-species coalescent (MSC) method with ASTRAL-Pro3 (Zhang & Mirarab, 2022). Based on partitioned multiple sequence alignments of single-copy HOGs, the ML analysis yielded a species tree with consistently high bootstrap support (Figure 1). Using gene trees from RAxML-ng, the MSC approach produced a fully supported species tree (posterior probability = 1 at all nodes; Supplementary Figure S8). Both methods resulted in largely congruent topologies, capturing major evolutionary lineages. However, both approaches revealed cross-genus nesting—*Pectis angustifolia* within *Porophyllum*, and *Coreopsis lanceolata* clustering outside its genus. Such cross-genus nesting has been previously reported for Asteraceae (Hansen et al., 2016; Huang et al., 2016b) and Brassicaceae (Huang et al., 2016a). Furthermore, the recovered intergeneric relationships and major clade topologies within the alliance align well with previous studies (Zhang et al., 2021).

**Figure 1:**
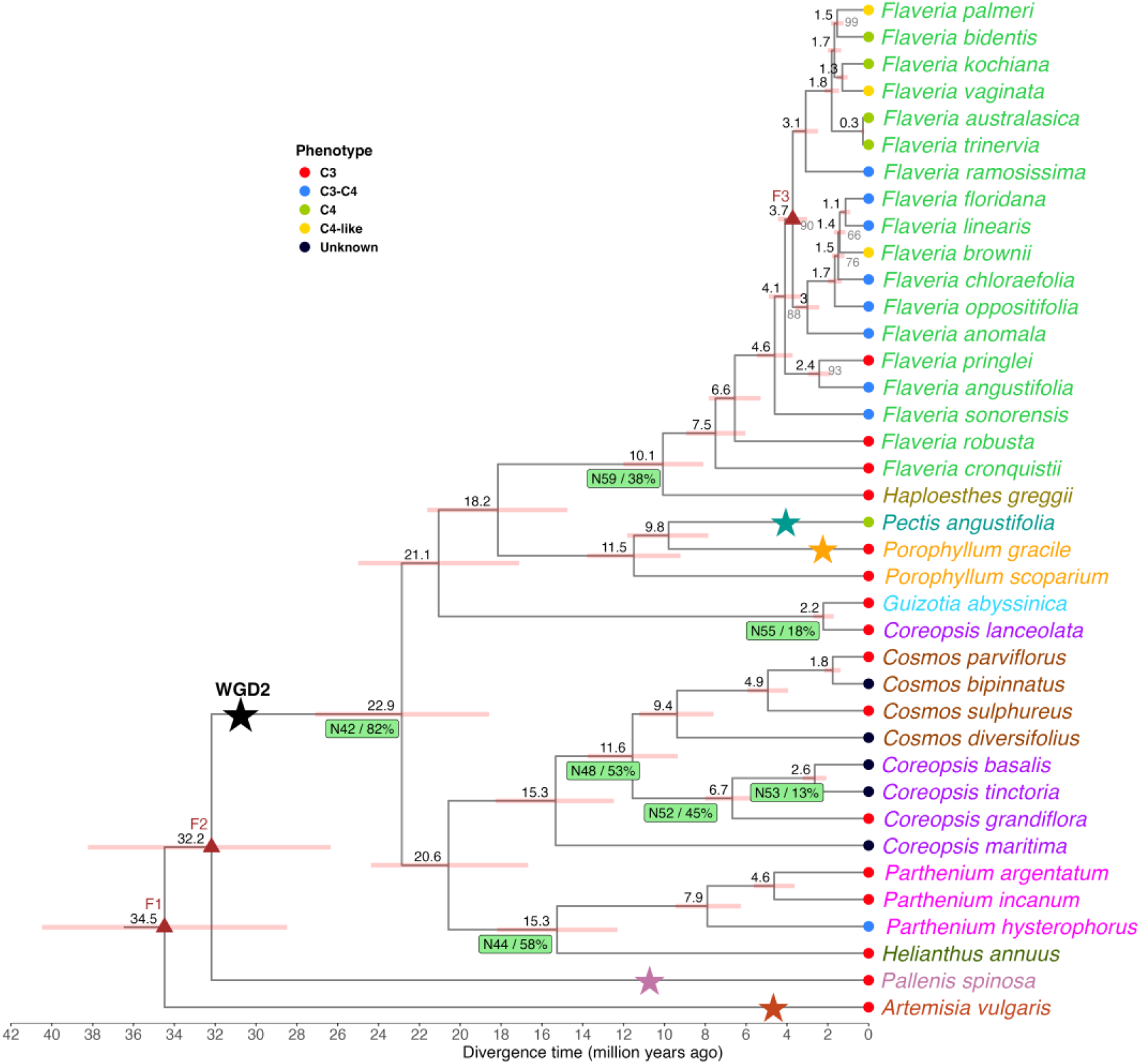
Species tree of 38 species belonging to *Heliantheae alliance*, *Anthemideae* and *Inuleae* tribes of *the Asteraceae* family. The focal species names are coloured according to their genus, while the tip-points are coloured according to their photosynthetic phenotype. All nodes showed a bootstrap of 100, otherwise mentioned in grey below the nodes. The internal nodes with brown triangles (F1, F2 and F3) indicate fossil calibrations obtained from the TimeTree database (http://www.timetree.org) used in speciation dating by MCMCtree in the PAML v4.10.7 package (Yang, 2007). The numbers in black above the nodes indicate the estimated divergence time in millions of years ago (MYA). The star in black indicates a shared whole-genome duplication (WGD) event called WGD-2, which occurred 30.84 million years ago (MYA) in the ancestors of the *Heliantheae alliance* species. The stars in green and orange colours indicate the additional and recent WGD events that occurred in respective species. The stars in purple and red colours indicate distinct WGD events that occurred in the respective species. The green rectangles above the nodes indicate WGD events shared by the descendant taxa. The first position indicates the node ID, and the second indicates the proportion of duplicate gene families shared by the descendant taxa in the rectangle.

Phylogenomics dating using MCMCtree (Yang, 2007) yielded divergence estimates broadly consistent with a recent phylotranscriptomic analysis (Zhang et al., 2021), placing the origin of the Heliantheae alliance at 32.2 million years ago (MYA). However, by leveraging 38 whole-genome sequences, we provide refined resolution for the most recent radiations, dating the Heliantheae-Coreopsideae split to 20.6 MYA and the divergence between the specialized Flaveria clades to 3.7 MYA (Figure 1). This refined timeline enables more precise associations between WGD-derived gene retention and the rapid evolution of C_4_ photosynthesis phenotypes.

### Identification of whole-genome duplications

Given the role of whole-genome duplications (WGD) in species diversification and evolutionary innovations in angiosperms (Soltis & Soltis, 2016; Van de Peer et al., 2017; One Thousand Plant Transcriptomes Initiative, 2019), we systematically identified WGD events across the studied 38 species to assess their prevalence and timing. We employed two complementary approaches to investigate WGDs: analysis of synonymous substitution rates (Ks) and gene tree-species tree reconciliation.

Using wgd v2 (Chen et al., 2024), we identified paralogous gene clusters and calculated Ks values for each paralogous gene pair within each species to identify Ks peaks corresponding to potential WGD events. All Heliantheae alliance species of our study exhibited a prominent Ks peak between 0.43 and 0.65 (mean: 0.51), suggesting a WGD event about 30.8 MYA (Figure 1 and Supplementary Figure S9). This peak aligns well with the previously reported ancient WGD-2 event (29 MYA) in the ancestor of *Helianthus annuus* (Badouin et al., 2017). In addition, we identified species-specific WGD events in *Pectis angustifolia* (Ks = 0.07; 4.24 MYA) and *Porophyllum gracile* (Ks = 0.04; 2.42 MYA), indicative of more recent duplications. Also, the outgroup species showed distinct WGD events: *Artemisia vulgaris* at Ks = 0.08 (4.85 MYA) and *Pallenis spinosa* at Ks = 0.18 (10.91 MYA) (Figure 1 and Supplementary Table S6).

To further validate these findings, we performed gene tree-species tree reconciliation using Tree2GD v2.5 (Chen et al., 2022). This analysis provided strong evidence for a shared ancient WGD event in the most recent common ancestor (MRCA) of the Heliantheae alliance (Node N42). A total of 82% of duplicated gene families were shared among descendant taxa, which was consistent with the Ks-based inference of the WGD-2 event (Figure 1 and Supplementary Table S7). The reconciliation approach also identified shared WGDs at the MRCAs of several genera, including *Helianthus* and *Parthenium* (Node N44), *Cosmos* and *Coreopsis* (Node N48), and *Flaveria* and *Haploesthes* (Node N59) (Figure 1). These results confirm that the observed duplications reflect large-scale genome duplications rather than small-scale gene duplications (Hakes et al., 2007).

Our results therefore highlight the dynamics of genome duplication in the evolutionary history of the Heliantheae alliance and its relatives.

### WGD-2-derived gene duplicates and their distributions

To assess the impact of the ancient whole-genome duplication event (WGD-2) on gene retention and functional conservation across species of the Heliantheae alliance, we analyzed more than 3.18 million paralog pairs (Supplementary Table S8). We focused on paralog pairs with Ks values of 0.41 – 0.68 (Supplementary Figure S10), which corresponds to a duplication time of ∼30.84 million years ago (Figure 1). A total of 17.6% of the total paralog pairs fulfilled this criterion. The proportion of WGD-2 duplicates varied widely across species (7.9% - 42.2%), with notable differences between absolute and relative counts, reflecting lineage-specific evolution (χ² = 121, p = 4.03 × 10⁻¹¹, Supplementary Figure S11A). Crucially, the genomic distribution and functional assignments of these retained pairs were highly non-random. On average, 60% of WGD-2 pairs were retained within the same Hierarchical Orthologous Groups, significantly deviating from random genomic allocation expectations (χ² = 183, p = 1.40 × 10⁻²¹, Supplementary Figure S11B and Supplementary Table S8). Furthermore, 85% of WGD-2 pairs shared the exact same MapMan BIN classifications, strongly rejecting the null hypothesis of neutral functional divergence (χ² = 79.3, p = 4.30 × 10⁻⁵, Supplementary Figure S11C and Supplementary Table S8). Together, these results suggest that a vast majority of WGD-2 duplicates actively maintained conserved biological roles and structural classifications, rather than undergoing rapid functional drift or decay following the duplication event.

### Phylogenetic association mapping of gene copy number variation with CO_2_ compensation points

Gene copy number variation (CNV) between species is a genetic polymorphism contributing significantly to phenotypic variability for adaptive evolution in plants and animals (Suryawanshi et al., 2016; DeBolt, 2010; Lye & Purugganan, 2019; Ishikawa et al., 2021; Wilhoit et al., 2025; Pajic et al., 2019; Aoyagi Blue et al., 2021). However, its role in diversifying carbon-concentrating mechanisms (CCMs) is largely unexplored. To address this gap, we associated CNV across 31 species from the Heliantheae alliance including two outgroups, to the CO₂ compensation point (CCP; Supplementary Table S9) as a quantitative proxy for CCM efficiency (Schlüter et al., 2023) using a phylogenetic association mapping (PAM) framework (Kiefer et al., 2019). This approach identified 742 high-confidence HOGs (Supplementary Table S10) and 451 MapMan BINs (Supplementary Table S11) with significant CNV-CCP associations (p < 0.05).

To gain insights into the functional relevance of the identified associations, we examined the biological roles of the MapMan BINs and HOGs identified through the PAM framework. Among the 451 MapMan BINs (containing ≥2 CNV states), twelve BINs correspond to key components of photosynthesis and photorespiration (Figure 2, Supplementary Table S12 and Figure S12). These include BINs related to both light-dependent and light-independent reactions: *Photosystem II LHC-II complex component* (LHCb5, BIN: 1.1.1.1.3), *Photosystem II assembly factor* (LTO1, BIN: 1.1.1.3.16), *Photosystem I LHC-I complex component* (LHCa3, BIN: 1.1.4.1.3), *ferredoxin electron carriers* (Fd1/2/3, BIN: 1.1.5.1.1), and *cyclic electron flow (PGRL1, BIN: 1.1.6.1.2)*—all integral to photophosphorylation. Additionally, the identification of *PTOX* (chlororespiration, BIN: 1.1.8.2), *BFA1* (ATP synthase assembly, BIN: 1.1.9.3.2), and enzymes of the Calvin Benson cycle, such as *GAPDH* (BIN: 1.2.3.1) and *fructose-1,6-bisphosphatase (BIN:* 1.2.6), reinforces the involvement of central photosynthetic machinery. Notably, two BINs involved in photorespiration (*glycine decarboxylase L-protein* (BIN: 1.3.4.3) and *hydroxypyruvate reductase HPR2/3, BIN:* 1.3.6.2) and one associated with C₄/CAM mechanisms (*carbonic anhydrase beta-type, BIN:* 1.4.1.2) were also identified. These findings are consistent with earlier reports that link evolutionary modifications in light-harvesting and Calvin cycle components to C₄ photosynthesis (Mallmann et al., 2014; Reyna-Llorens & Hibberd, 2017).

**Figure 2:**
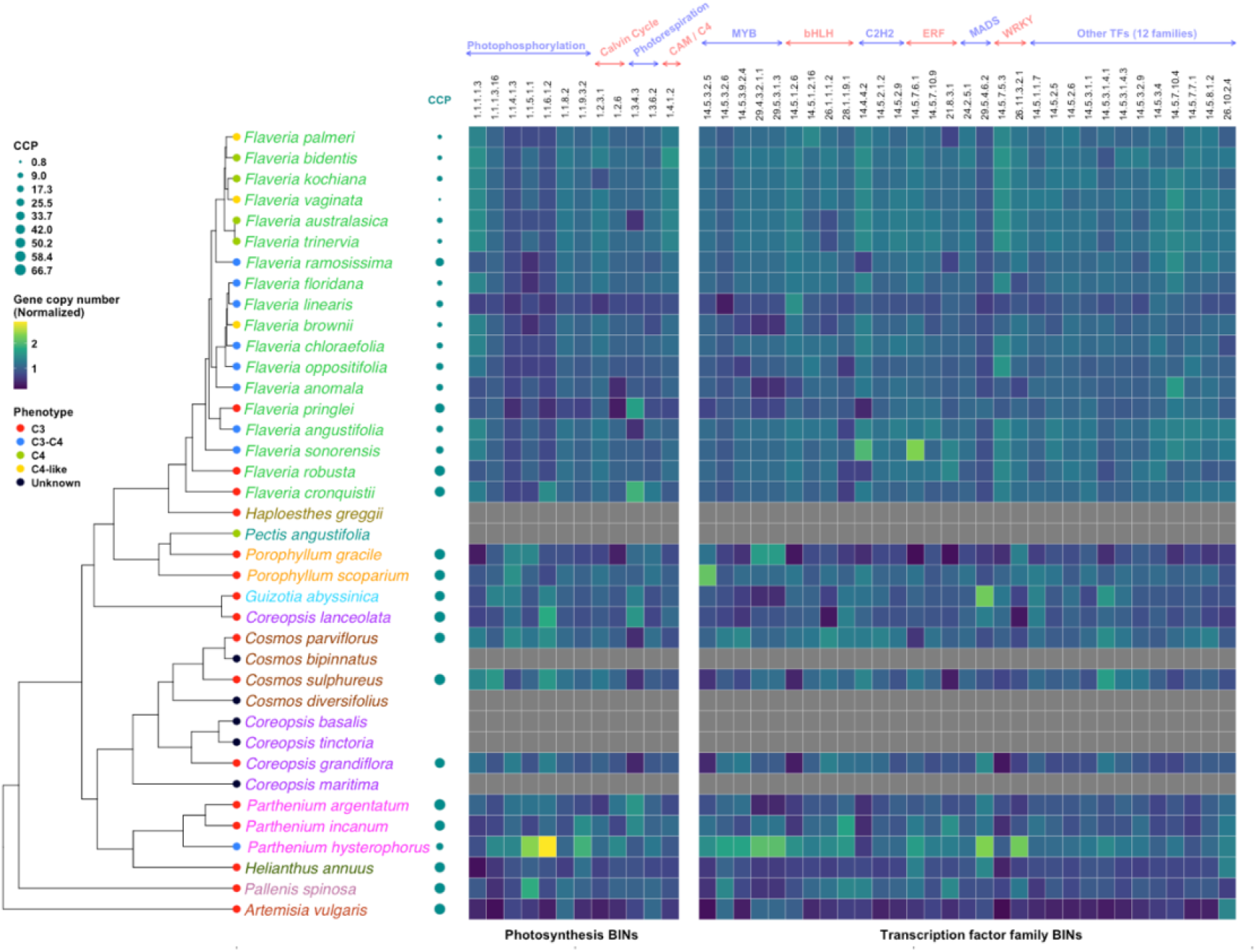
Phylogenomic distribution of CO_2_ compensation point (CCP)-associated MapMan BINs for photosynthesis and transcription factor families. This integrated analysis depicts the distribution of gene copy number variations (CNVs) across the species tree (derived from Figure 1) for 43 CCP-associated functional categories. The heatmap is partitioned into two vertical blocks: 12 MapMan BINs related to **photosynthesis** (left) and 31 BINs related to **transcription factor families** (right). Heatmap cell values represent normalized gene copy numbers. Species-specific physiological traits are displayed alongside the tree, including photosynthetic types (colored points) and CCP values (proportional circle sizes). Grey cells in the heatmap indicate taxa excluded from the PAM analysis due to the unavailability of CCP values.

In parallel, 31 transcription factor-related BINs were significantly associated with CCP values (Figure 2, Supplementary Table S13 and Figure S13). These spanned multiple TF superfamilies, including MYB, bHLH, C2H2, ERF, MADS, WRKY, AHL, BEL, bZIP, DREB2, GeBP, GRAS, GRF, HD-ZIP, HSF, KNOX, LOB and TCP families. For the MYB-domain transcription factors this is consistent with recent findings that link MYB TFs to chloroplast biogenesis and photosynthetic regulation (Frangedakis et al., 2024). Similarly, the identification of bHLH, bZIP, and WRKY transcription factors aligned with evidence that these TFs play key roles in regulating photorespiration (Aroca et al., 2023).

### Non-linear CNV-trait associations and effect sizes

To determine whether CNV-CCP association reflected gradual dosage effects or a threshold-dependent architecture, we visually examined the linear relationships between CNV and CCP for each significant BIN. CNV-CCP distributions revealed no consistent monotonic trends of linear relationships across all associated BINs. These results indicate that the detected phylogenetic associations are not explained by simple additive or linear scaling of gene copy number.

Given the absence of consistent linear relationships, we tested whether these associations followed a non-linear, threshold-based architecture by calculating the systematic effect size (η^2^) and genomic expansion magnitude (Cohen’s *d*) across all 451 BINs (see Materials and Methods). These analyses indicate that the transition from C_3_ to C_4_ followed a tipping-point framework (Figure 3A) similar to that described for speciation (Nosil et al., 2017; Dakos et al., 2019).

**Figure 3:**
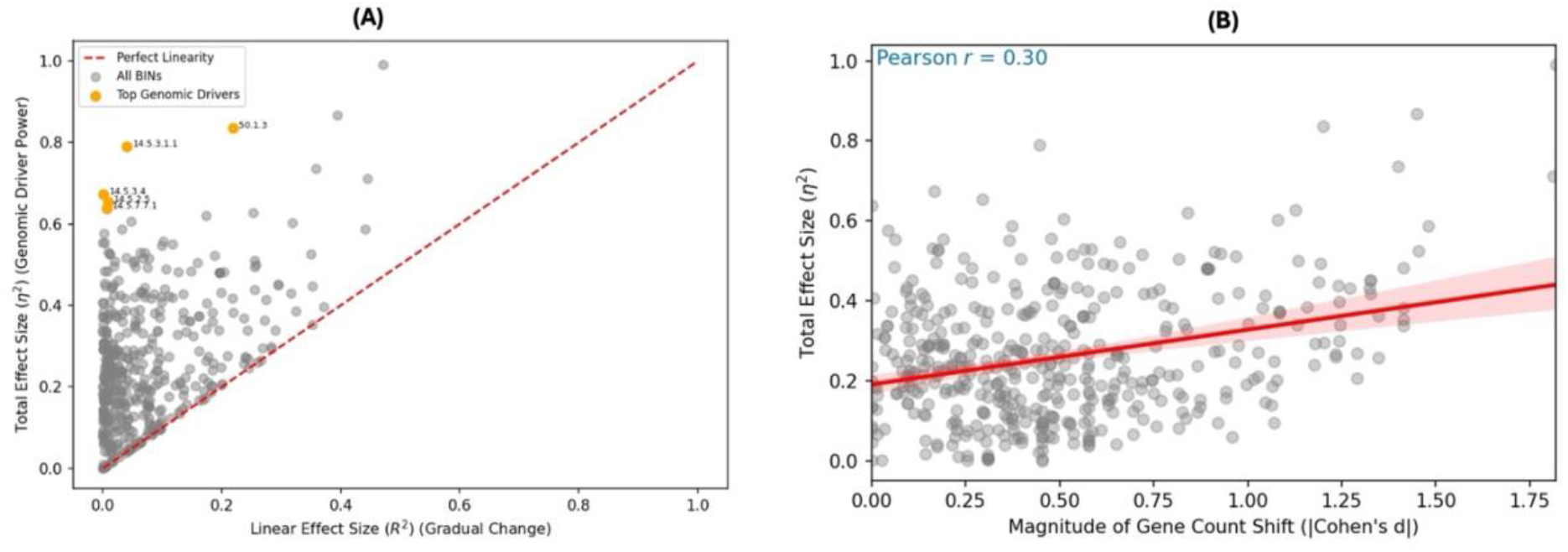
Genomic tipping points characterize the C_3_-to-C_4_ physiological transition. **(A) Decoupling of linearity and explanatory power.** Total effect sizes (η^2^) plotted against linear R^2^ for 451 MapMan BINs significantly associated with CCP values. The five most extreme ‘tipping points’ (highest η^2^ - R^2^ gap) are highlighted: transcription factor BINs: 14.5.3.1.1 (HD-ZIP), 14.5.3.4 (HSF), 14.5.2.5 (LOB), 14.5.7.7.1 (TCP), and 50.1.3 (Oxidoreductase). Their position in the upper-left quadrant indicates that they are massive drivers of CCP variation that cannot be explained by linear models. **(B) Magnitude-Impact Scaling.** Correlation between the scale of genomic change (*d*) and driver power (η^2^). The significant positive trend confirms that larger shifts in gene copy number are the primary engine behind high-magnitude physiological shifts.

Across the significant BINs, the total explanatory power of gene dosage on CCP varied substantially. The median η^2^ was 0.24, and the maximum reached 0.99, even though linear correlations (R^2^) were negligible (Figure 3A; Supplementary Table S14). We then identified the top five primary drivers (BINs) with the highest tipping point magnitude (defined as the non-linear surplus, η^2^ -R^2^). These included transcription factor categories 14.5.3.1.1 (HD-ZIP), 14.5.3.4 (HSF), 14.5.2.5 (LOB), 14.5.7.7.1 (TCP), and BIN 50.1.3 (Oxidoreductase). Together, these BINs explained between 63% and 83% of the total variance in CCP, while each showed only a weak linear correlation (R^2^ <0.22; Figure 3A, orange color circles). We also detected a positive correlation (r = 0.3) between the magnitude of gene counts shift (Cohen’s *d*) and physiological impact (Figure 3B). Although moderate in strength, this relationship suggests that larger genomic expansions are more likely to generate the biological thresholds associated with CCM evolution.

### Validation of the identified phylogenetic associations

Because of the observed low contribution of each individual HOG or BIN to variation in the CCP values, we did not validate them on an individual basis using functional genetic approaches but as a group using genomic prediction models. The results suggested that the BINs that were identified using the above-mentioned PAM approach resulted in a significantly higher prediction performance (Friedman test, *p* = 0.006 6 for *R*² and *p* = 0.004 for RMSE; Nemenyi post hoc test, *p* < 0.05) than all BINs, whereas PAM-derived HOGs did not differ significantly from all HOGs (Figure 4A). Specifically, across all examined validation schemes, PAM-derived BINs and PAM-derived HOGs showed lower error (RMSE = 14.7 ppm, 15.0 ppm) and the higher out-of-sample *R*² (0.475, 0.456), outperforming models that include all BINs (15.6 ppm; *R*² 0.408) or HOGs (15.3 ppm; *R*² 0.426) (Figure 4A). In measured hold-out tests, these four feature sets were statistically indistinguishable (*p*>0.05) with best performance around RMSE = 11.3 ppm and *R*² = 0.58, indicating minimal overfitting (Figure 4B). Furthermore, we confirmed the robustness of these associations under distribution shift scenarios using both independent external data (literature-based validation) and artificially perturbed datasets with swapped species assignments (swap-resampled validation). Across these tests, PAM-derived BINs consistently maintained a performance advantage over full BIN sets (e.g., *R*² = 0.489 vs. *R*² = 0.352 for swap-resampling; Figure 4B).

**Figure 4:**
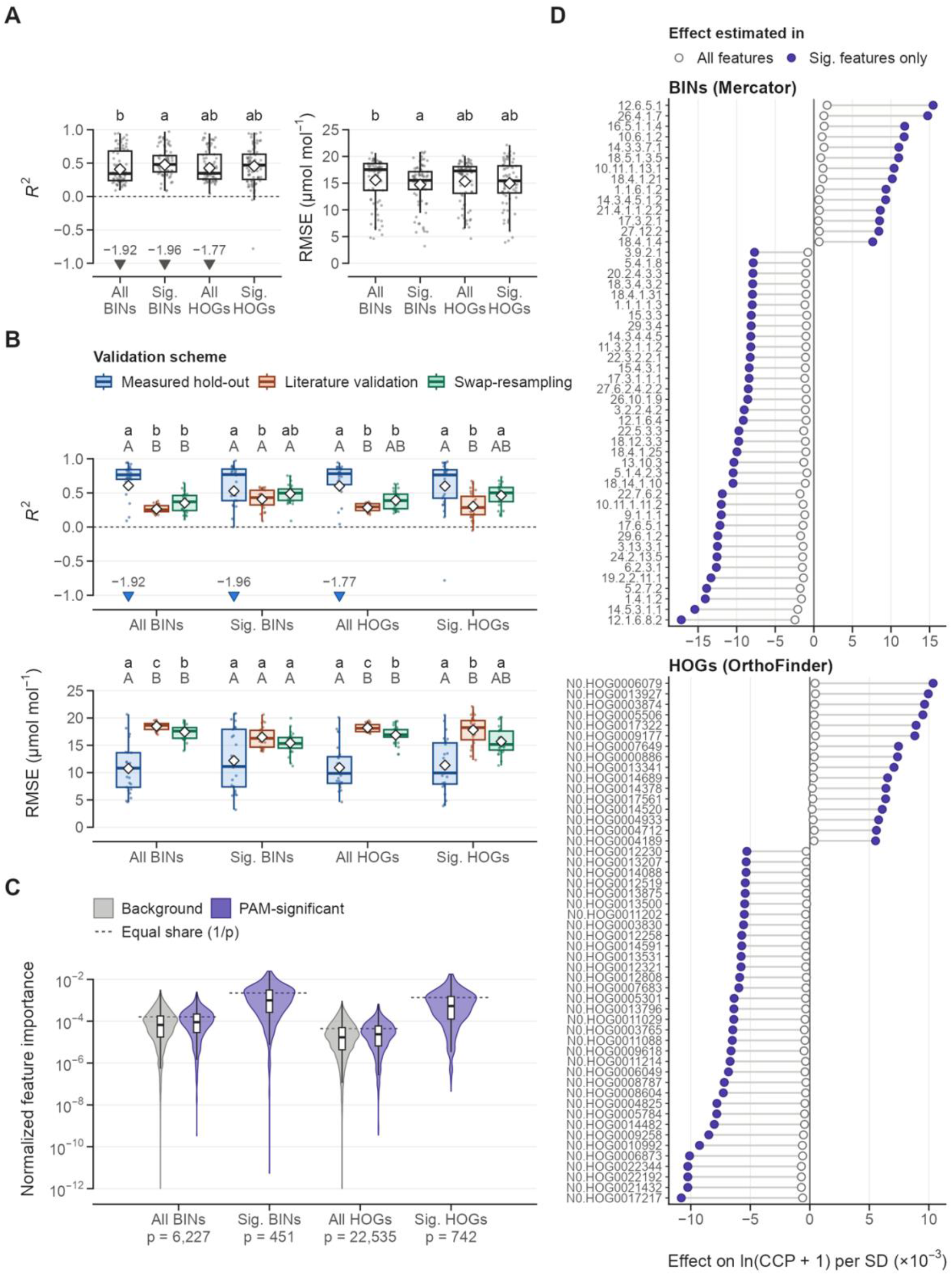
A) Distribution of genomic prediction performance metrics (out-of-sample R² and root mean square error RMSE) across feature set, and B) validation schemes. C) Normalized squared rrBLUP effects for PAM-significant and background gene-family predictors. D) Top 50 CCP predictors across feature sets (HOGs and BINs): ranked effects and overlap.

To identify the most relevant gene families among the set of PAM associations, we ranked them based on the absolute standardized rrBLUP effect size from the genomic prediction models (Figure 4D). The top 50 PAM-derived BINs included functions relevant to CCP biology, such as photosynthesis light reactions, chlorophyll and tetrapyrrole metabolism, photorespiration and glyoxylate pathways, Calvin-Benson cycle, mitochondrial energy metabolism, redox and ROS homeostasis, and metabolite transport (Figure 4D and Supplementary Table S15).

### Contribution of whole-genome duplication-2 to CCP-associated MapMan BINs’ functional categories

After having established that copy number variation in specific MapMan BINs strongly correlates with variation in CCP and the validation procedure, we next sought to determine the evolutionary origin of these expansions. Specifically, we examined whether the ancient whole-genome duplication (WGD-2) event served as the primary genomic substrate for these adaptive gene expansions. In order to do so, we quantified the proportion of WGD-2–derived genes within functionally annotated MapMan BINs across all species.

Out of the 451 BINs for which CNV was significantly associated with CCP, 303 BINs (67.18%) contained at least 10% of their genes derived from WGD-2 (Supplementary Table S16). To determine whether this genomic retention was non-stochastic, we performed a genome-wide hypergeometric enrichment test against the background of all 6,472 annotated MapMan BINs. This analysis confirmed that the functional modules linked to CCP variation are non-randomly enriched for WGD-2 retentions (p = 2.84 x 10^-46^), establishing a broad, systemic influence of WGD-2 on CNV at the BIN-level. In addition, the same approach indicated that the WGD-2 retention in BINs of the structural and regulatory machinery of the carbon-concentrating mechanism are significantly (p = 9.12 x 10^-4^) over-represented (Figure 5 and Supplementary Table S12). Furthermore, the genomic legacy of WGD-2 extended deeply into upstream regulatory networks. Twenty-one out of the 31 transcription factor (TF) BINs (67.7%) crossed the 10% WGD-2 retention threshold, with proportions ranging from 12.87% to 87.30% (Figure 5 and Supplementary Table S13). The hypergeometric enrichment test indicated that this duplication-driven expansion within TF regulatory pathways is also highly significant compared to the background genome (p = 2.33 x 10^-4^).

**Figure 5:**
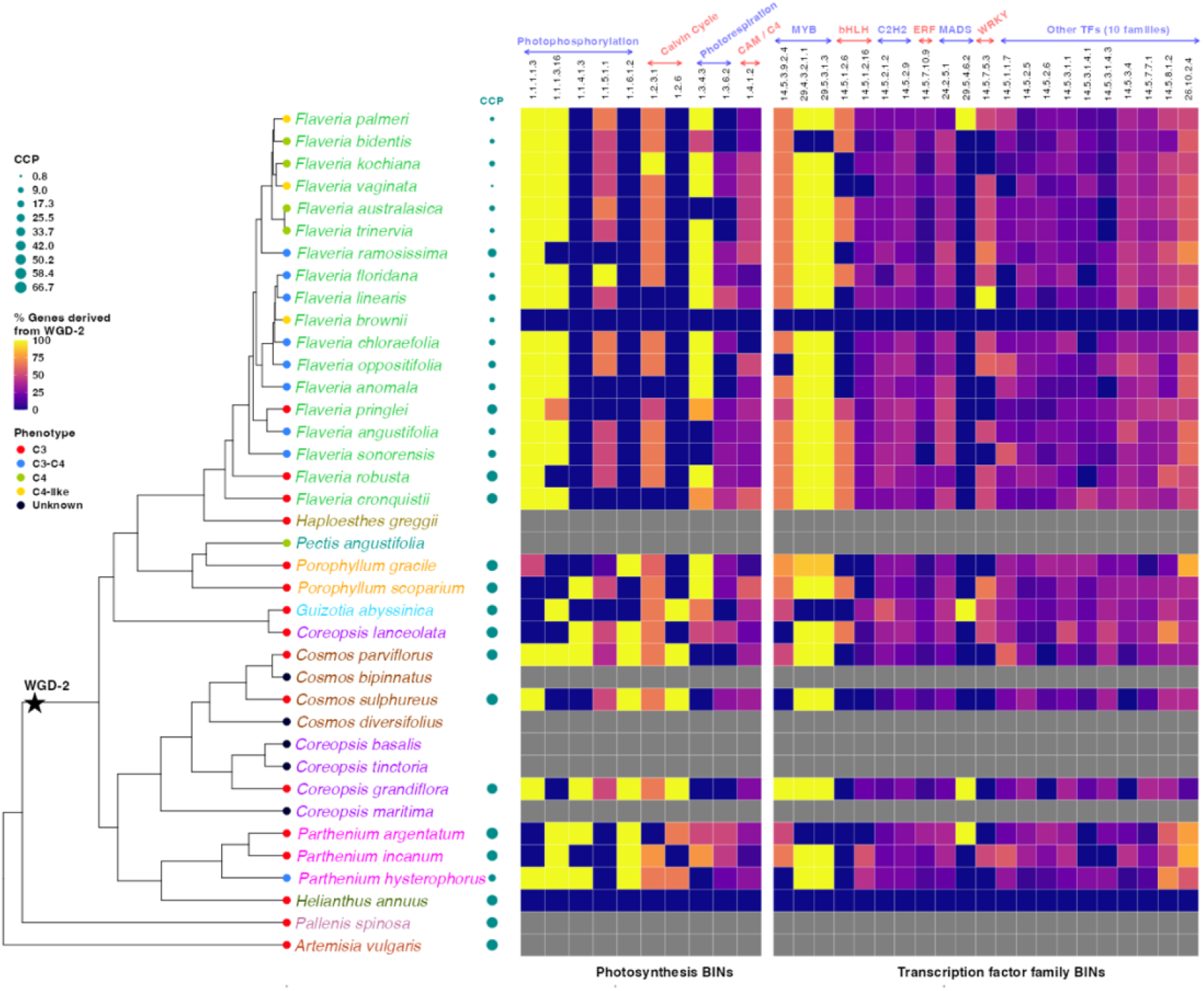
Genomic contribution of the WGD-2 event to CCP-associated MapMan BINs. This integrated figure illustrates the proportion of CCP-associated CNVs originating from the WGD-2 event, mapped across the Asteraceae phylogeny (based on the tree topology in Figure 1). The heatmap displays the retention patterns for functional categories associated to the CO_2_ compensation point (CCP), partitioned into **photosynthesis-related BINs** (left) and **transcription factor families** (right). The heatmap gradient represents the percentage of the total CNVs within each BIN that is traceable to the WGD-2 event. Gray cells represent taxa excluded from the PAM and WGD analyses due to missing physiological CCP data or the absence of discernible WGD-2 genomic peak.

## Discussion

The repeated, independent emergence of carbon-concentrating mechanisms (CCMs) represents one of the most significant physiological transitions in plant evolution (Sage, 2004; Sage et al., 2011, 2012). The Asteraceae—particularly the Heliantheae alliance—provide a unique natural system to investigate this phenomenon. This is due to its exceptional photosynthetic diversity and deep phylogenetic divergence allowing to bridge the gap between ancient polyploidization events and present-day physiological phenotypes.

### High-quality genome assemblies and evolutionary context as foundations for trait mapping

Because none or only assemblies of limited-quality were previously available (cf. Taniguchi et al., 2021), we generated new, high-quality reference resources for these lineages as part of our 36 new assemblies. This improvement is central to our study, as the reliability of interspecies CNV-phenotype associations depends strictly on genome assembly quality. The assemblies presented in our study were generated using diverse sequencing technologies, including short-read, linked-read and long-read platforms. Despite this heterogeneity, all assemblies exhibit a high contiguity and completeness (Supplementary Figures S1-S4) and met the critical quality threshold of an N50 greater than the average gene length (Supplementary Table S3), which is a prerequisite for accurate gene and paralog identification (Yandell & Ence, 2012). Furthermore, BUSCO completeness scores remained consistently high and displayed minimal inter-species variation (Supplementary Figure S2), indicating that the gene space is sufficiently captured across all assemblies to enable robust, comparable CNV estimation. Furthermore, the quality of the assemblies is comparable to, or exceeds, existing Asteraceae assemblies (Taniguchi et al., 2021) and is similar to assemblies from well-studied Brassicaceae species (Kiefer et al., 2019; Guerreiro et al., 2023).

Annotation bias is a common challenge in multi-platform comparative studies. To address this issue, we applied a uniform, centralized annotation pipeline across all species. This pipeline incorporated the deep learning-based tool Helixer (Holst et al., 2026) and InterProScan (Jones et al., 2014). This consistency was essential to ensure that variation in gene counts reflect true biological copy number variation (CNV) and is not due to technical artifacts or inconsistent gene calling. Therewith our PAM analysis was grounded on high-quality, uniformly processed genomic data providing a robust comparative framework for all following analyses.

### WGD-2: A shared genomic innovation with functional legacy

Whole genome duplications are widely recognized as engines of plant diversification and trait innovation (Jiao et al., 2011; Soltis & Soltis, 2016; Van de Peer et al., 2017; Landis et al., 2018; One Thousand Plant Transcriptomes Initiative, 2019). They also create opportunities for new traits to emerge. Despite this, their effects on complex traits like photosynthesis are less clear. Past studies often focused on isolated examples or a small set of pathway-specific genes (Ludwig, 2011; Schulze et al., 2013; Bianconi et al., 2018; Huang et al., 2021; Hoang et al., 2023), but our study provides a systematic analysis of how this process shaped complex photosynthetic phenotypes.

In this study, we identified a conserved WGD-2 event (K_s_ = 0.43–0.65) that is shared across all Heliantheae alliance species, placing the duplication at their most recent common ancestor (Figure 1). This genomic redundancy precedes therewith the diversification of all C_3_-C_4_ and C_4_ lineages within the Heliantheae alliance (Badouin et al., 2017). Our observation that the MapMan BINs associated with CO_2_ compensation point (CCP) variation are significantly enriched for WGD-2 derived genes illustrate that the WGD-2 event is not merely resulting in ancestral “baggage” but has a functional relevance. Our results therefore directly support the “pre-conditioning” hypothesis of Monson (2003): the WGD-2 event expanded the genomic repertoire of the Heliantheae ancestor, providing a “modular” set of duplicated genes that later acted as evolutionary hubs for the assembly of complex CCM phenotypes (Supplementary Figure S11). These observations are giving an evolutionary explanation to the results of earlier studies which revealed the duplication of a narrow set of core photosynthesis genes (Ludwig, 2011; Schulze et al., 2013; Taniguchi et al., 2021; Hoang et al., 2023) while we move beyond and reveal how WGD-2 provided the genomic raw material for the systemic anatomical and metabolic rewiring necessary for CCM evolution which is discussed in the following for different pathways.

### WGD-2-driven expansion of photosynthetic and regulatory modules supports CCM evolution

#### 1. Metabolic specialization and energy flux

While the ancient WGD-2 event established the foundational genomic scaffold for these pathways, subsequent lineage-specific duplications provided the “fine-tuning” necessary for tissue-specific optimization. Together, these events expanded critical metabolic nodes, such as β-carbonic anhydrase (β-CA), which is essential for the initial conversion of CO_2_ to HCO ^-^(Hatch & Burnell, 1990). The retention of multiple WGD-2 derived β-CA copies, supplemented by lineage-specific expansions (Figure 5; Supplementary Table S12), suggests that selection in the Heliantheae favored both higher enzyme capacity and precise control of carbon flow.

Furthermore, the co-retention of duplicated Calvin-Benson cycle enzymes (GAPDH, FBPase) and light-harvesting components (LHCb5, LHCa3) likely enabled subfunctionalization among these gene copies (Figure 5; Supplementary Table S12). Such partitioning would help balance RuBP regeneration and light capture between mesophyll and bundle sheath cells. It meets the higher ATP demands of CCM-active species and reduces stress from reactive oxygen that arises with increased carbon flux (Rochaix, 2011).

#### 2. Regulatory and anatomical innovation

A clear example for the refinement of the duplication space provided by the WGD-2 event by lineage-specific gene family expansions is the SCARECROW (SCR) gene family (GRAS transcription factors); WGD-2 derived SCR copies likely allowed ancestral root endodermis networks to be co-opted for leaf patterning, facilitating the characteristic Kranz anatomy (Slewinski et al., 2012; Hughes et al., 2019; Shaar-Moshe & Brady, 2023).

This regulatory rewiring extends to bHLH and MYB transcription factor families. MYB proteins likely promote CCM evolution by modulating gene networks involved in chloroplast biogenesis, enhancing the anatomical and metabolic differentiation required for efficient glycine shuttling in C_3_-C_4_ intermediates. Such control may have facilitated the transition from C_3_ to C_3_-C_4_ photosynthesis by optimizing organelle development (Frangedakis et al., 2024; Dickinson et al., 2025). Furthermore, the synergistic binding of duplicated bHLH transcription factors to C_4_ promoters, such as NADP-ME (Borba et al., 2018), provides one possible mechanism for functional enhancement. More gene copies and stronger promoter binding probably raised local CO_2_ around Rubisco, helping to reduce photorespiration.

The concurrent expansion of both core metabolic BINs and regulatory transcription factor modules, driven by the synergy of WGD-2 and subsequent lineage-specific duplications, appears to underlie the variation in CO_2_ compensation points (CCP) across the Heliantheae alliance. Furthermore, our findings demonstrate that CCM evolution is not driven by isolated enzymes, but by the coordinated, modular expansion of entire metabolic and regulatory landscapes.

Because of the observed low contribution of each individual HOG or BIN to variation in the CCP values, we did not try to validate them on an individual basis using functional genetic approaches but as a group using genomic prediction models (Dos Santos Sá et al., 2026). In detail, we selected the BINs for which the CNV was significantly associated with CCP values in the PAM. Those were combined and used as a group to predict CCM values of a so-called validation set. We observed that genomic prediction with phylogenetically informed BINs improved cross-species accuracy for CCP relative to unfiltered feature sets. This indicated that the evolutionary-based grouping helps retain conserved, biologically meaningful signal and that the previously described associated CNV have biological relevance.

### Non-linear Architecture Underlies CNV-CCP Associations

The linear effect sizes (R^2^) for individual BIN copy numbers are often modest, but the total explanatory power (η^2^) is substantially higher (Figure 3; Supplementary Table S14). This difference suggests that gene dosage effects on CCP are not additive. Rather, these effects appear shaped by non-linear genetic architectures, such as evolutionary thresholds or tipping points (Nosil et al., 2017; Dakos et al., 2019). This observation can be explained by the fact that duplicated genes do not act alone. They work within complex regulatory networks that include epistatic interactions and feedback loops (Kaste et al., 2024). Epigenetic regulation and context dependent gene expression may further disconnect raw copy numbers from the resulting traits (Hennacy & Jonikas, 2020; Cummins, 2021). In summary, whole-genome duplication gives the raw material for evolutionary innovation. However, translating these expansions into functional physiological adaptations requires complex, multi-layered mechanisms, and non-linear mechanisms.

## Conclusion

By integrating high-quality genome assemblies with experimental measurements across 35 species spanning C_3_, C_3_-C_4_ intermediate, and C_4_ photosynthetic types, we established a genome-wide interspecies framework linking carbon-concentrating mechanism (CCM) evolution to genomic features within the Heliantheae alliance.

Crucially, we identified a shared whole-genome duplication event dated to approximately 30 million years ago (Figure 1), which established a foundational genomic scaffold that subsequent lineage-specific duplications refined to drive contemporary trait variation. Phylogenetic association mapping revealed systematic gene copy number variations across critical functional categories (Figure 2), with photosynthetic associations enriched in functional modules containing at least 10% WGD-2-derived genes.

Rather than involving isolated loci, these expansions are distributed across coordinated biochemical and regulatory BINs—encompassing both metabolic enzymes and upstream regulatory transcription factors. This pattern indicates that CCM evolution relied on the retention and diversification of duplicated functional modules.

Therewith, this work provides a refined genomic blueprint for enhancing photosynthetic efficiency in major Asteraceae crops, including sunflower. Instead of focusing on single enzymes, breeders could aim to increase the copy number of these regulatory genes, through genome editing or by inducing polyploidy and selecting the best-performing lines, as a way to improve photosynthesis in crops such as sunflower.

## Materials and Methods

### Species selection and physiological characterization

We selected 35 Asteraceae species from the Heliantheae alliance and closely related tribes (Supplementary Table S1) spanning the full photosynthetic continuum, from C_3_ to C_4_ types. All species were grown under controlled conditions (24°C/21°C day/night; 16h photoperiod). These standardized settings ensured that physiological measurements were directly comparable across species. To quantify the carbon concentrating mechanism (CCM) efficiency, we measured CO_2_ compensation points (CCP) using infrared gas exchange measurements (LI-COR 6800). CCP values were derived from A/c_i_ curves under high light (1500 µmol m^-2^ s^-1^) and a controlled vapor pressure deficit of 1.5 kPa. The detailed growth conditions and gas exchange protocols are described in the Supplementary Information.

### Multi-omics sequencing and genome assembly

To establish a high-resolution genomic framework, we generated *de novo* genome assemblies for 35 species. High-molecular-weight DNA was extracted using an optimized CTAB-based protocol specifically adapted for long-read library preparation (Arseneau et al., 2017; Xin & Chen, 2012). Genome sequencing was realized using a combination of short-read, linked-read, and long-read sequencing technologies (Supplementary Table S1 and S2). To support gene model prediction and quantify gene activity, we performed multi-organ transcriptomic sequencing (RNA-seq) for 27 species and direct-RNA sequencing for 8 more species. Genome assemblies were generated using assemblers, such as Flye (Kolmogorov et al., 2019), Supernova (Weisenfeld et al., 2017), and DeNovoMAGIC (NRGene, Nes Ziona, Israel). Assembly quality was evaluated based on contiguity and completeness via BUSCO analysis (Seppey et al., 2019). Detailed descriptions of DNA and RNA extraction, library preparation, and assembly parameters are provided in the Supplementary Information.

### Phylogenomics and divergence dating

Gene families, defined as hierarchical orthogroups (HOGs), were inferred across 38 species using OrthoFinder v2.5.5 (Emms & Kelly, 2019). Species phylogeny was constructed using two complementary approaches. First, we performed maximum-likelihood analysis of concatenated single-copy orthologs using RAxML-ng (Kozlov et al., 2019). Second, we applied a multi-species coalescent framework using ASTRAL-Pro3 (Zhang & Mirarab, 2022). Divergence times were estimated using a Bayesian relaxed-clock model implemented in MCMCtree (Yang, 2007). This analysis was calibrated with three fossil-based reference points. Partition models, bootstrap parameters, and calibration priors are detailed in the Supplementary Information.

### Identification of whole-genome duplication (WGD)

Paralogous gene clusters were identified across species. For each paralog pair, synonymous substitution rates (Ks) were calculated to detect historical polyploidy events. WGD-associated Ks peaks were detected using kernel density estimation. Confidence intervals (90%) were obtained through resampling. To distinguish shared from lineage-specific duplication events, gene duplication nodes were mapped onto the species tree using Tree2GD (Chen et al., 2022). A detailed description of the Ks filtering pipeline and reconciliation criteria is provided in the Supplementary Information.

### Phylogenetic association mapping (PAM)

To link genomic architecture with physiological traits, we applied the phylogenetic association mapping framework of (Kiefer et al., 2019). This approach accounts for evolutionary relatedness via a variance-covariance matrix derived from the species phylogeny. Normalized gene copy numbers (HOGs and MapMan BINs) were regressed against CCP values across 31 species. Predictive performance was evaluated using ridge-regression best linear unbiased prediction (rrBLUP; Endelman, 2011). Model robustness was assessed through repeated stratified cross-validation. Although we performed phylogenetic association mapping using 31 species for which CCP measurements were available, we excluded two species (A. vulgaris and P. spinosa), for which the WGD-2 peak was absent, from this analysis. The mathematical derivation of the normalization formulas, the structure of the predictive models, and the cross-validation schemes are elaborated in the Supplementary Information.

### Quantifying effect sizes and evolutionary tipping points

To characterize the non-linear relationship between CNV and CCP, we calculated two complementary statistics across all 451 BINs: the systematic effect size (η^2^) and the genomic expansion magnitude (Cohen’s *d*). η^2^ quantifies the proportion of total variation in CCP explained by differences in copy number states and provides an overall measure of the explanatory power of a given MapMan BIN with respect to CCP. To quantify the magnitude of genomic shift between photosynthetic states, we additionally calculated Cohen’s *d*. Together, these metrics allowed the detection of potential evolutionary tipping points (Nosil et al., 2017; Dakos et al., 2019).

## Data availability

Sequencing data for all 35 species have been deposited in the NCBI BioProject database (https://www.ncbi.nlm.nih.gov/bioproject) under accession number PRJNA1260212. Sequence data for *Flaveria bidentis* was separately obtained from BioProject PRJNA1259992. Genome assemblies and annotations have been archived on FigShare and are available at https://doi.org/10.6084/m9.figshare.c.8797566.

## Code availability

All scripts used in this study are available on GitHub at https://github.com/vsbonthala/Gain4Crops.

## Author contributions

BS conceptualized the study and acquired funding. BVS performed the genome assembly, annotation, and all phylogenomic analyses—including species tree construction, WGD identification, association mapping, and quantification of WGD-derived CNVs—and drafted the manuscript with input from all authors. SK managed plant growth and tissue collection, US conducted the CCP measurements, and KMK validated the association mapping through genomic predictions.

## Acknowledgments

We gratefully acknowledge the following institutions and genebanks for generously providing seed and germplasm materials essential for this study: the USDA-ARS National Arid Land Plant Genetic Resources Unit (Parlier, CA, USA), the USDA-ARS Western Regional Plant Introduction Station at Washington State University (Pullman, WA, USA), the Leibniz Institute of Plant Genetics and Crop Plant Research (IPK, Gatersleben, Germany), the Royal Botanic Gardens, Kew (UK), and the Botanical Garden of Heinrich Heine University Düsseldorf (Germany). The authors acknowledge the computational infrastructure and support provided by the Center for Information and Media Technology at Heinrich Heine University Düsseldorf and the German Network for Bioinformatics Infrastructure (de.NBI, https://www.denbi.de/) that contributed to the research results reported within this study. The authors gratefully acknowledge the computing time provided to them on the high-performance computers [Noctua2] at the NHR Center PC2. These are funded by the Federal Ministry of Education and Research and the state governments participating on the basis of the resolutions of the GWK for the national high-performance computing at universities (www.nhr-verein.de/unsere-partner).

We thank Prof. Tomas Morosinotto (University of Padova, Italy) and Dr. Giovanni Finazzi (French Alternative Energies and Atomic Energy Commission, CEA, France) for their help with the photosynthetic phenotyping.

## Supplementar Figures

**Figure S1: Assembly quality and genomic annotations.** The bar plot displays the assembly quality and genomic annotations for the newly assembled 35 species. Assembly size in gigabases (A), Assembly contiguity (log10 N50 (Kb)) (B), Percentage of assembly completeness (BUSCO scores) (C), Repetitive sequences (D), Genic regions (E), and Assembly gaps (F) of the assembly of each species. Green bars represent species with assemblies (reference-quality) generated using short-read and mate-pair libraries, followed by scaffolding with 10X linked reads using DeNovoMAGIC (NRGene, Nes Ziona, Israel). Purple bars represent species assembled with a single stLFR library using Supernova v2.1.1 (Weisenfeld et al., 2017), and orange bars represent species assembled with an Oxford Nanopore long-read library using Flye v2.9 (Kolmogorovet al., 2019) followed by scaffolded with a single stLFR linked-read library. The species are sorted based on the descending order of N50 values.

**Figure S2: BUSCO completeness across 35 species assemblies.** The horizontally stacked bar plot displays the BUSCO (Benchmarking Universal Single-Copy Orthologs) completeness scores for the newly assembled 35 species, assessing the quality of their genome assemblies. Each bar represents a species and is colour-coded to reflect the proportion of complete (light blue), duplicated (blue), fragmented (yellow), and missing (red) BUSCO genes.

**Figure S3: Distribution of transposable elements across species.** The horizontally stacked bar plot shows the composition of the top ten transposable elements (TE) groups across 35 species. Each bar represents a species, with the width of each coloured segment corresponding to the percentage of the assembly occupied by a specific TE group. The top ten TE groups are colour-coded and include elements such as Copia, Gypsy, the long-terminal repeats (LTRs), Helitrons and Mutators, among others.

**Figure S4: Comparative genomic features across species assemblies.** The horizontal bar plot consists of seven subplots, each representing a different genomic feature for the newly assembled 35 species and *Flaveria brownii* (Cyan colour bar). The features include (A) the total number of genes, (B) the mean gene length in base pairs, the percentage of genes with UniProt descriptions (C), PFAM protein domains (D), Gene Ontology terms (E), Expressed (F), and MapMan Ontology BINs (G). We considered genes with transcripts per million (TPM) of 0.5 or greater as expressed or those with TPM values greater than 0 in the case of gene expression measured using direct RNA sequencing data (as indicated by “*”). The bar with “**” indicates that the publicly available RNA-seq data was used to calculate the expression. The bars are colour-coded, and species are sorted as in Supplementary Figure S1.

**Figure S5: Correlation between genome size, transposable elements (TEs), and gene content across 35 species.** The two plots display the relationships between genome size and two genomic features: (A) a solid positive correlation (R²: 0.995) is observed between the increase in transposable elements (TEs) and the increase in genome size, indicating that TE expansion is a significant contributor to genome size variation. (B) A moderate correlation (R²: 0.544) is observed between gene content and genome size, suggesting that gene content plays a lesser role in genome size changes than TEs. In both plots, species are grouped and colour-coded by genus, highlighting the trends across taxonomic groups.

**Figure S6: Phylogenetic Hierarchical Orthogroups (HOGs) across species.** This figure displays an overview of HOGs and their distribution across multiple species. It provides insights into the percentage of genes assigned, HOG counts, and the types of HOGs identified by OrthoFinder v2.5.5 **(Emms & Kelly, 2019)**. A) Percentage of genes in each species assigned to HOGs. B) Total number of HOGs identified for each species. C) Distribution of HOG types across species: This stacked bar plot represents the proportion of different HOG types in each species. We categorised HOGs into three groups - HOGs containing all species, HOGs containing more than one species but not all, and species-specific HOGs, i.e., HOGs with just one species. D) Distribution of genes in HOG types: Similar to subplot 3, this plot focuses on the distribution of genes in HOGs. It shows the percentage of genes associated with HOGs that are either present in all species, shared across some species, or species-specific.

**Figure S7: Pairwise heatmap of orthologous gene sharing between 38 species.** This heatmap represents the percentage of genes shared as orthologs between each pair of species, as determined by OrthoFinder v2.5.5 (Emms & Kelly, 2019). Each cell in the heatmap shows the percentage of orthologous genes shared between two species, ranging from low (red) to high (blue). Higher percentages suggest closer evolutionary relationships or similar gene content, while lower percentages indicate more divergent genomes or gene content.

**Figure S8: Estimation of Species tree using ASTRAL-Pro3.** We reconstructed a species tree using ASTRAL-Pro3 (Zhang & Mirarab, 2022), incorporating gene trees constructed by RAxML-ng (Kozlov et al., 2019) of 5,618 Hierarchical Orthogroups (HOGs) that contain all 38 species. Quartet scores at each internal node are represented by pie charts, illustrating the relative proportions of the three possible quartet relationships: Q1 (Concordance), Q2 (Alternative), and Q3 (Conflict). Each pie chart visually depicts the consensus among the gene trees. All internal nodes have a posterior probability (PPP) of 1, indicating strong support for the branching structure.]

**Figure S9: Ks distribution density with predicted peaks for 38 Species.** This figure presents the density distributions of synonymous substitution rates (Ks) computed using wgd v2 (Chen et al., 2024) across 38 species, with each subplot representing a single species. We identified the predicted peaks of whole-genome duplication events using Gaussian Mixture Modelling (GMM). To assess the robustness of these predictions, we performed resampling 1,000 times, each time selecting half of the Ks values, to calculate 90% confidence intervals for each peak. Subplots are colour-coded according to the genus of each species. The density plots highlight the distribution of Ks values, indicating possible whole-genome duplication (WGD) or other duplication events across species.

**Figure S10: Representation of whole-genome duplication-2 in Ks distribution.** The WGD peak range corresponds to the minimum and maximum Ks of peaks identified across species. The red and purple lines indicate the combined lower and upper bounds of 90% confidence intervals (CI) of the WGD peak across all focal species.

**Figure S11: Duplicated gene pairs derived from whole-genome duplication-2 (WGD-2).** The bar plot displays (A) the percentage of duplicated genes derived from whole-genome duplication-2 (Ks = 0.41-0.68). The Chi-square test revealed a statistically significant difference in the proportion of duplicated gene pairs derived from WGD-2 across species (χ² = 121, p-value: 4.03 × 10⁻¹¹). (B) The proportion of duplicated gene pairs derived from WGD-2 sharing the same or different hierarchical orthologous groups (HOG; χ² = 183, p = 1.40 × 10⁻²¹). (C) The proportion of duplicated gene pairs derived from WGD-2 sharing the same or different MapMan BIN (χ² = 79.3, p = 4.30 × 10⁻⁵). Here, *Artemisia vulgaris* and *Pallenis spinosa* were excluded from the analysis because both lack a WGD-2 peak within the Ks range of 0.41-0.68.

**Figure S12: Association between Gene Copy Numbers of BINs related to photosynthesis and Carbon Compensation Point (CCP).** Boxplot showing the distribution of carbon compensation point (CCP) values (Y-axis) across species grouped by gene copy number variation (X-axis). Each boxplot represents one MapMan BIN associated with CCP.

**Figure S13: Association between Gene Copy Numbers of BINs related to transcription factors and Carbon Compensation Point (CCP).** Boxplot showing the distribution of carbon compensation point (CCP) values (Y-axis) across species grouped by gene copy number variation (X-axis). Each boxplot represents one MapMan BIN associated with CCP.

