## Supplementary Information for "Ancestral polyploidy and non-linear dosage shifts define the photosynthetic transition in the Heliantheae alliance"

### Species selection and physiological characterization

#### Selection of species and plant growth

We collected seeds from various gene banks for 35 Asteraceae species that have been previously categorized as performing $C_3$, $C_3$-$C_4$, $C_4$-like, and $C_4$ photosynthesis (Supplementary Table S1). Seeds were germinated and plantlets grown in growth chambers using the following conditions: 7% sand and 93% soil mixture; 16h light (24°C, 70% humidity) and 8h dark (21°C, 70% humidity). Plants were placed in darkness for 48h before collecting tissue samples. We collected tissues from 1-4 organs, such as flowers, leaves, stems, and roots, in liquid nitrogen for later sequencing genomes and transcriptomes (Supplementary Table S1).

#### Measuring CO2 compensation points

*Flaveria* species *F. cronquistii*, *F. chloraefolia*, *F. linearis*, *F. angustifolia*, *F. oppositifolia*, *F. sonorensis*, *F. kochiana*, and *F. vaginata* were grown from cuttings in Floraton 1 soil mixture (Floraton 1, Floraguard, Germany). For all other species, seeds were vapour sterilized by incubation in an exicator together with a fresh mixture of 100 ml of 13% Na-hypochloride and 3 ml of 37% HCl for 2 hours. The sterilized seeds were then germinated on plates containing 0.22% (w/v) Murashige Skoog medium, 50 mM MES at pH 5.7, and 0.8% (w/v) Agar. After 7 to 10 days, the seedlings were transferred to Floraton 1 soil. All plants were cultivated in a greenhouse with a 16-hour day and 8-hour night cycle. Natural light conditions in the greenhouse were supplemented with Valoya RX600 LED lamps, enabling the plants to receive a light intensity of between 250 and 800 µmol m-2 s-1. Minimum temperatures were adjusted to 21°C during the night and 24°C during the day.

Gas exchange was measured on the youngest fully expanded rosette leaf about 6 to 10 weeks after sowing or transplantation and before the onset of flowering. The settings of the LI6800 (LI-COR Corporate, USA) were as follows: flow rate of 300 µmol s-1, fan speed of 100,000 rpm, light intensity of 1500 µmol m-2 s-1, leaf temperature of 25°C and VPD of 1.5 kPa. After adjusting the leaves to the conditions in the leaf chamber, A-ci curves were measured by setting reference CO2 concentrations at 400, 300, 200, 100, 80, 60, 40, 20, and 10 ppm. The CO2 compensation points (CCPs) were calculated using a minimum of four data points in the linear range close to the interception with the Ci axis.

### Multi-omics sequencing, genome assembly, and annotation

#### DNA extraction and sequencing

High-molecular-weight (HMW) DNA was extracted using a modified protocol based on Arseneau et al. (2017) and Xin and Chen, (2012). For each sample, four 50 mg aliquots of leaf tissue were homogenized in 1 mL of preheated (60°C) CTAB lysis buffer (100 mM Tris-HCl, 20 mM EDTA, 2% CTAB, 1.2 M NaCl, 0.1% ß-mercaptoethanol). Following incubation at 60°C for 1 hour (300 rpm, inverting every 15 min), samples were cooled to room temperature, mixed with 1 mL chloroform, and centrifuged at 3,000g for 15 min at room temperature.

The upper aqueous phase (650 µl) was transferred to a new tube and mixed with 1,300 µl of diluted CTAB buffer (100 mM Tris-HCl, 20 mM EDTA, 2% CTAB). After a second incubation (60°C, 1 h, 600 rpm) and centrifugation (16,000g, 5 min), the supernatant was discarded. To remove excess CTAB, the pellet was soaked in 1 mL of 80% ethanol for 30 min at room temperature. After final centrifugation (16,000g, 15 min), the pellet was resuspended in 25 µl high-salt TE buffer (10 mM Tris, 2 mM EDTA, 1 M NaCl), and the four batches per sample were pooled (100 µl total).

RNA was degraded by adding 1 µl RNase A (100 mg/mL, Qiagen) at 37°C for 1 hour. DNA was purified using AMPure XP beads (1:1 ratio), washed three times with 80% ethanol, and air-dried. The beads were resuspended in 50 µl Buffer AE (Qiagen). DNA was allowed to solubilize for 1 hour at room temperature and stored overnight at 4°C prior to bead removal.

Purified samples for 26 species were dispatched to BGI Genomics (Shenzhen, China) for library preparation and high-throughput sequencing. The remaining ten samples were dispatched to NRGene, Nes Ziona, Israel for library preparation and high-throughput sequencing.

#### RNA extraction and sequencing

Total RNA was isolated using the RNeasy Plant Mini Kit (Qiagen) following the manufacturer's protocol. RNA integrity was assessed via capillary electrophoresis (e.g., Agilent Bioanalyzer 2100), ensuring a RNA Integrity Number (RIN) > 7.0 prior to shipment. Purified samples were dispatched to BGI Genomics (Shenzhen, China) and Heinrich Heine University, Düsseldorf, Germany for library preparation and high-throughput sequencing.

**Whole-genome sequencing**

We sequenced the genomes of 26 species using the single-tube long fragment read (stLFR) linked-read technology (Wang et al., 2019) at BGI, China, and one species with Oxford Nanopore long-read sequencing technology at GTL, Heinrich Heine University, Düsseldorf, Germany. We sequenced nine species using 10X linked-read and Illumina whole-genome sequencing technologies to create reference-quality genome assemblies at NRGene, Nes Ziona, Israel. We generated 10X linked reads, shotgun sequences with 440 and 700 base inserts, and three different-length mate-pair libraries: 2-4 Kb, 5-7 Kb, and 8-10 Kb (Supplementary Table S1, Supplementary Table S2).

**Transcriptome sequencing**

We sequenced the transcriptomes of at least one organ, such as flower, leaf, stem and root, for 27 species using Illumina short-read technology at BGI, China. In addition, we sequenced the transcriptomes of the RNA extracted from pooled organs of eight species using Oxford Nanopore direct RNA sequencing technology at GTL, Heinrich Heine University, Düsseldorf, Germany (Supplementary Table S1).

**Estimation of genome sizes**

We estimated genome sizes for all newly sequenced species using a K-mer-based approach. We counted K-mers of lengths 19 to 63 using Jellyfish v2.3.1 (Marçais & Kingsford, 2011), followed by an estimated genome size calculation using GenomeScope (Vurture et al., 2017) and GenomeScope2 (Ranallo-Benavidez et al., 2020).

**NRGene DeNovoMAGIC assemblies**

The shotgun and mate-pair sequence libraries were used for *de novo* genome assembly using the DeNovoMAGIC (NRGene, Nes Ziona, Israel), a DeBruijn-graph-based genome assembler to generate reference-quality assemblies. The raw Chromium 10X data was utilised to phase polyploidy/heterozygosity, support scaffolds validation and further elongation of the phased scaffolds.

**Nanopore long-read assembly**

The raw sequencing reads of *Flaveria palmeri* were filtered to remove short (<5 kb) and/or reads with q < 9 using NanoFilt v2.8 (De Coster et al., 2018). Filtered reads were used for *de novo* assembly using Flye v2.9 (Kolmogorov et al., 2019) with default parameters, except that the number of polishing iterations was set to three. The assembled reads were further polished using filtered reads using three rounds using Racon v1.4.3 (Vaser et al., 2017) and one round with Medaka v1.6.0 (https://github.com/nanoporetech/medaka). To remove redundant contigs resulting from residual heterozygosity and individual haplotype assembly, we ran purge_haplotigs v1.1.2 (Roach et al., 2018) on the assembly using default parameters. To further enhance the contiguity of the assembly, we performed scaffolding of the assembled sequences using ARCS (Yeo et al., 2018) + LINKS (Warren et al., 2015) pipeline (https://github.com/bcgsc/arcs) using stLFR linked reads.

**stLFR linked-read assemblies**

We generated stLFR linked-read assemblies using Supernova v2.1.1 (Weisenfeld et al., 2017) with default parameters using the stlfr2supernova pipeline (https://github.com/BGI-Qingdao/stlfr2supernova_pipeline). Briefly, this pipeline involves filtering low-quality reads (Q < 10), PCR duplicates and adapters using SOAPfilter v2.2 (Luo et al., 2012), converting stLFR-linked reads into 10X-linked reads, and finally running Supernova assembler. To remove redundant scaffolds resulting from residual heterozygosity and individual haplotype assembly, we ran purge_haplotigs v1.1.2 (Roach et al., 2018) on Supernova's assemblies using default parameters.

**Assessing the quality of the assemblies**

We conducted a quality assessment of the assemblies, evaluating the contiguity of the assembled sequences using the N50 metric. We also assessed the completeness of the assemblies for capturing conserved single-copy orthologous genes using Benchmarking Universal Single-Copy Orthologs (BUSCO) v5.5.0 against the "embryophyta_odb10" lineage dataset (Seppey et al., 2019).

**Genome annotation**

Whole-genome transposable elements (TE) annotation was performed for all newly assembled species using the Extensive *de-novo* TE Annotator (EDTA) v2.0.0 with default parameters (Ou et al., 2019). To ensure efficient TE annotation, we fed a repetitive elements library of *Viridiplantae* obtained from RepBase (Bao et al., 2015) to EDTA.

We performed gene annotations for all newly assembled species and *Flaveria brownii* by feeding the genome assemblies to Helixer v0.3 (Holst et al., 2026), a deep learning-based gene annotation tool trained on a "land_plant" dataset to annotate protein-coding genes. Furthermore, we performed functional annotation for the predicted genes using the standalone version of InterProScan v5.57-90.0 (Jones et al., 2014) to assign functional descriptions, including InterProScan descriptions, Pfam protein domains, and Gene Ontology terms. We performed MapMan Bin annotation (Schwacke et al., 2019) for predicted genes using the Mercator4 tool (Lohse et al., 2014). In addition, we assigned gene descriptions by performing BLASTP (Altschul et al., 1990) with default parameters against the UniProt protein database downloaded (on 29/August/2022) from https://www.uniprot.org. Finally, we removed gene models producing protein sequences with <50 amino acids and gene models related to transposable elements using a curated list of Pfam protein domain IDs (Jayakodi et al., 2020).

#### Gene expression analysis

We mapped the RNA-seq data of each organ against the mRNA sequences of the respective species using Kallisto v0.48.0 (Bray et al., 2016) to quantify the expression in terms of transcripts per million (TPM). We defined genes with a TPM value greater than 0.5 as expressed in the respective tissue.

The obtained reads were subjected to quality filtering to quantify direct-RNA sequence reads. We used NanoFilt v2.8.0 (De Coster et al., 2018) to remove reads with length <150 bases and quality with q <10, followed by trimming reads with adapters and splitting reads with internal adapters using Porechop v0.2.4 (Wick et al., 2017). Finally, we estimated expression abundance (TPM) using high-quality direct RNA sequencing reads with NanoCount v1 (Gleeson et al., 2022). We defined genes with a TPM value greater than 0 as expressed.

### Phylogenomics and divergence dating

#### Computing gene families

We analyzed predicted proteomes from 38 species. Gene models were newly predicted for 36 species in the current study, while annotated proteomes for two additional species were obtained from published genome resources, including *Flaveria bidentis* (Sun et al., 2025) and *Helianthus annuus* (Badouin et al., 2017). The combined dataset comprised 1,339,739 protein sequences.

All proteomes were fed to OrthoFinder v2.5.5 (Emms & Kelly, 2019) using the default parameters to compute phylogenetic hierarchical orthogroups (HOGs) or gene families.

**Estimation of the species tree**

We estimated species trees using two approaches. In the first approach, we extracted the protein sequences of 76 single-copy HOGs and aligned them using MAFFT v7.520 (Katoh & Standley, 2013). We back-translated the protein alignment into nucleotide alignment using PAL2NAL v14 (Suyama et al., 2016) and concatenated the individual alignments into a single large matrix of alignment. PartitionFinder (Lanfear et al., 2017) was used to identify the optimal partitions for the concatenated alignment and to select the most suitable evolutionary models for each partition during species tree reconstruction. Using RAxML-ng v1.2.2 (Kozlov et al., 2019), we estimated a species tree by inputting the identified partitions, their corresponding models, and the concatenated alignment with 1000 bootstrap replicates.

In the second approach, we performed a multiple sequence alignment for each HOG, comprising all 38 species, at the protein sequence level using MAFFT v7.520 (Katoh & Standley, 2013) and then back-translated the alignment into a nucleotide alignment using PAL2NAL v14 (Suyama et al., 2016). The alignments were fed to RAxML-ng (Kozlov et al., 2019) to independently reconstruct phylogenetic trees for each HOG using the "GTRGAMMA" model and 200 bootstrap replicates. Finally, we fed the gene trees to ASTRAL-Pro3 (Zhang & Mirarab, 2022) to compute a species tree.

**Speciation dating**

Based on the species tree obtained from RAxML-ng (the first approach mentioned above), we performed speciation dating among 38 species using the program MCMCtree in the PAML v4.10.7 package (Yang, 2007). After completing a burn-in of 10,000 iterations, we ran the Markov chain Monte Carlo (MCMC) 100,000 times. We calibrated the divergence time estimation using three reference points: 1) the divergence time between *Anthemideae* (*Artemisia vulgaris*) and *Inuleae* (*Pallenis spinosa*) is 28.5 - 40.6 million years ago (MYA), 2) the divergence time between *Inuleae* (*Pallenis spinosa*) and *Heliantheae alliance* (*Helianthus annuus*) is 23.2 - 38.8 MYA, and 3) the divergence time between clade A (*Flaveria brownii*) and clade B (*Flaveria bidentis*) of Flaveria genus is 2.2 - 9.6 MYA, according to data from TimeTree (http://www.timetree.org).

### Identification of whole-genome duplication (WGD)

#### Identification of whole-genome duplications

We used the DMD program from WGD v2 (Chen et al., 2024) with default settings to identify paralogous gene clusters. Then, we used the KSD program of WGD v2 (Chen et al., 2024) with default settings to calculate the number of synonymous substitutions per synonymous site (Ks) values for each gene pair of paralogous gene clusters. We averaged Ks values at each node within the clusters to avoid redundant estimates. We applied Kernel density estimation (KDE) to the Ks distributions between 0.001 and 2 to detect peaks corresponding to whole-genome duplications (WGDs), followed by resampling to calculate 90% confidence intervals. Specifically, we performed 1,000 resamplings, each time selecting half of the Ks values, and repeated the KDE to obtain the confidence interval. These estimates were converted into time estimates using the Asterid substitution rate ($r = 8.25 \times 10^{-9}$ substitutions per synonymous site per year), following the formula divergence date = Ks/(2 × r), as reported by Badouin et al. (2017). The final estimate for the date of the WGD event is an average value of individual estimates. In comparison, we selected the confidence interval to include all individually estimated confidence intervals.

#### Identification of shared whole-genome duplications using phylogenetic reconciliation

We used the reconciling gene trees with the species tree method to identify WGD events shared by two or more species and place them on the species tree. We selected gene trees from all HOGs generated by RAxML-ng (Kozlov et al., 2019). We fed these phylogenetic trees and the species tree to Tree2GD v2.5 (Chen et al., 2022) with the default parameters to estimate the gene duplications (GD). We retained nodes with a GD of 300 or greater for further analysis. We considered WGD events to have occurred at a node according to any of the following criteria: 1) GD >= 2000, of which the proportion of (AB)(AB) type duplications are over 10%; 2) GD >= 1000, of which the proportion of (AB)(AB) type duplications are over 20%; 3) GD >= 500, of which the proportion of (AB)(AB) type duplications are over 30%; 4) GD >= 300, of which the proportion of (AB)(AB) type duplications are over 50%.

### Phylogenetic association mapping (PAM)

#### Phylogenetic association mapping

We applied the phylogenetic association mapping (PAM) method developed by Kiefer et al. (2019) to link the genotype (gene copy numbers of HOGs and MapMan BINs) against the phenotype (CO2 Compensation Points). To compare the copy number of MapMan BINs and HOGs between species, we normalized the copy number of genes within each MapMan BIN and HOG into ratios in the focal species by the total number of genes in the species to account for the species with a large number of genes would have a large number of genes in respective MapMan BIN and HOG using the formula mentioned in Aoyagi Blue et al. (2021). The phenotype (CCP) was used as the response variable, a fixed effect was used for the copy number of MapMan BINs and HOGs, and a random effect for species. We included 27 species for the CCP values have been measured in this study. Additionally, we included four species for which the CCP values were obtained from the literature (Supplementary Table S9).

The identified associations were evaluated as a group by evaluating the cross-species predictive performance of gene-family profiles for CCP. We fitted ridge-regression best linear unbiased prediction (rrBLUP) models using normalized HOG and BIN count matrices and their PAM-significant subsets as predictors. For each predictor set, we performed 25 independent runs. In each run, four measured species were randomly selected as hold-out test set, and the four species with literature-derived values were excluded from training. Models were trained on the remaining measured species using repeated stratified 5-fold cross-validation (10 repeats, 4:1 train:test ratio) and used to predict the held-out measured species and literature-only species. To assess sensitivity of predictions to train-test set composition, we carried out swap-resampling in each run by exchanging one measured and one literature species between training and prediction pools (5 swaps/run). Prediction performance was summarized using root mean square error (RMSE) and out-of-sample $R^2 = 1 - \frac{\Sigma(y-\hat{y})^2}{\Sigma(y-\bar{y}_{test})^2}$, where $\bar{y}_{test}$ is the mean of the test set (Campbell and Thompson, 2008). All analyses were performed in R using the rrBLUP package (Endelman, 2011).

**Non-linear CNV-trait associations and effect sizes**

Unlike linear correlation coefficients, $\eta^2$ captures both linear and non-linear components of association. This is achieved by partitioning total variance ($SS_{total}$) into variance between copy-number groups ($SS_{between}$) and variance within groups.
