## Supplementary Figures for "Ancestral polyploidy and non-linear dosage shifts define the photosynthetic transition in the Heliantheae alliance": Figure.S2.pdf

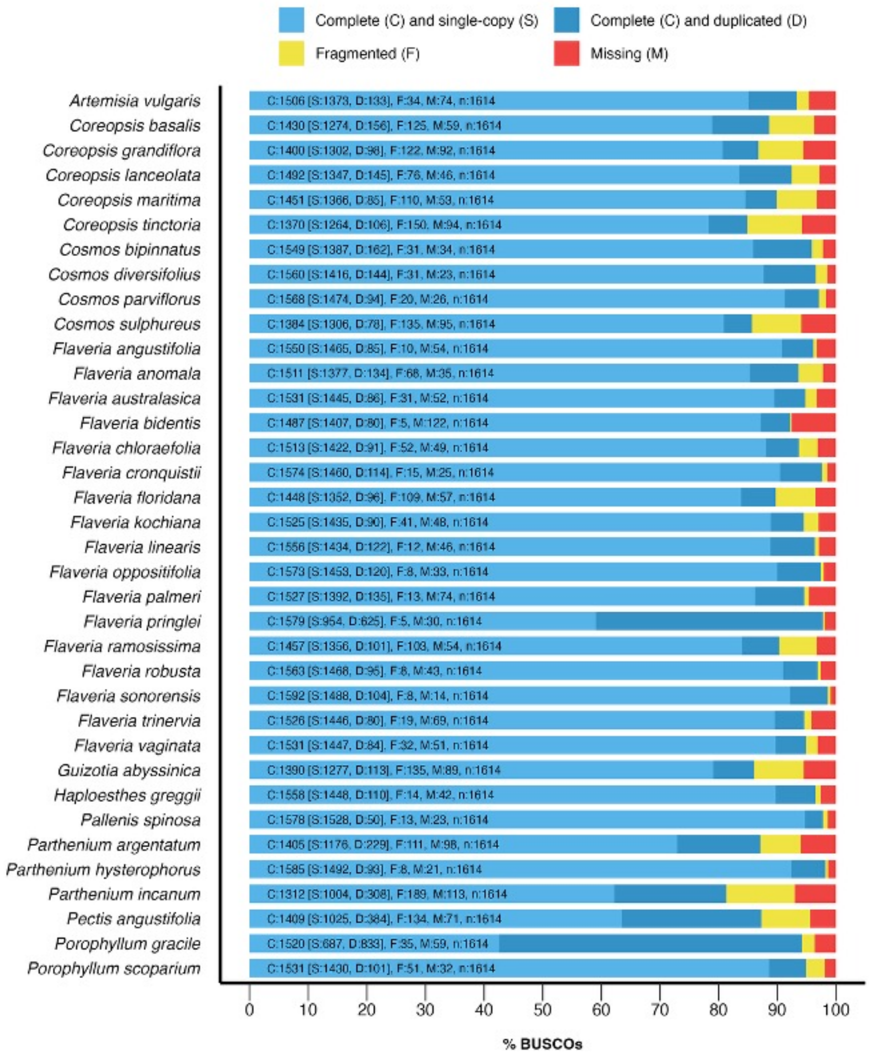
Complete (C) and single-copy (S)
Complete (C) and duplicated (D)
Fragmented (F)
Missing (M)
Artemisia vulgaris
C:1506 [S:1373, D:133], F:34, M:74, n:1614
Coreopsis basalis
C:1430 [S:1274, D:156], F:125, M:59, n:1614
Coreopsis grandiflora
C:1400 [S:1302, D:98], F:122, M:92, n:1614
Coreopsis lanceolata
C:1492 [S:1347, D:145], F:76, M:46, n:1614
Coreopsis maritima
C:1451 [S:1366, D:85], F:110, M:53, n:1614
Coreopsis tinctoria
C:1370 [S:1264, D:106], F:150, M:94, n:1614
Cosmos bipinnatus
C:1549 [S:1387, D:162], F:31, M:34, n:1614
Cosmos diversifolius
C:1560 [S:1416, D:144], F:31, M:23, n:1614
Cosmos parviflorus
C:1568 [S:1474, D:94], F:20, M:26, n:1614
Cosmos sulphureus
C:1384 [S:1306, D:78], F:135, M:95, n:1614
Flaveria angustifolia
C:1550 [S:1465, D:85], F:10, M:54, n:1614
Flaveria anomala
C:1511 [S:1377, D:134], F:68, M:35, n:1614
Flaveria australasica
C:1531 [S:1445, D:86], F:31, M:52, n:1614
Flaveria bidentis
C:1487 [S:1407, D:80], F:5, M:122, n:1614
Flaveria chloraefolia
C:1513 [S:1422, D:91], F:52, M:49, n:1614
Flaveria cronquistii
C:1574 [S:1460, D:114], F:15, M:25, n:1614
Flaveria floridana
C:1448 [S:1352, D:96], F:109, M:57, n:1614
Flaveria kochiana
C:1525 [S:1435, D:90], F:41, M:48, n:1614
Flaveria linearis
C:1556 [S:1434, D:122], F:12, M:46, n:1614
Flaveria oppositifolia
C:1573 [S:1453, D:120], F:8, M:33, n:1614
Flaveria palmeri
C:1527 [S:1392, D:135], F:13, M:74, n:1614
Flaveria pringlei
C:1579 [S:954, D:625], F:5, M:30, n:1614
Flaveria ramosissima
C:1457 [S:1356, D:101], F:103, M:54, n:1614
Flaveria robusta
C:1563 [S:1468, D:95], F:8, M:43, n:1614
Flaveria sonorensis
C:1592 [S:1488, D:104], F:8, M:14, n:1614
Flaveria trinervia
C:1526 [S:1446, D:80], F:19, M:69, n:1614
Flaveria vaginata
C:1531 [S:1447, D:84], F:32, M:51, n:1614
Guizotia abyssinica
C:1390 [S:1277, D:113], F:135, M:89, n:1614
Haploesthes greggii
C:1558 [S:1448, D:110], F:14, M:42, n:1614
Pallenis spinosa
C:1578 [S:1528, D:50], F:13, M:23, n:1614
Parthenium argentatum
C:1405 [S:1176, D:229], F:111, M:98, n:1614
Parthenium hysterophorus
C:1585 [S:1492, D:93], F:8, M:21, n:1614
Parthenium incanum
C:1312 [S:1004, D:308], F:189, M:113, n:1614
Pectis angustifolia
C:1409 [S:1025, D:384], F:134, M:71, n:1614
Porophyllum gracile
C:1520 [S:687, D:833], F:35, M:59, n:1614
Porophyllum scoparium
C:1531 [S:1430, D:101], F:51, M:32, n:1614
0
10
20
30
40
50
60
70
80
90
100
% BUSCOs
