## Supplementary Figures for "Ancestral polyploidy and non-linear dosage shifts define the photosynthetic transition in the Heliantheae alliance": Figure.S3.pdf

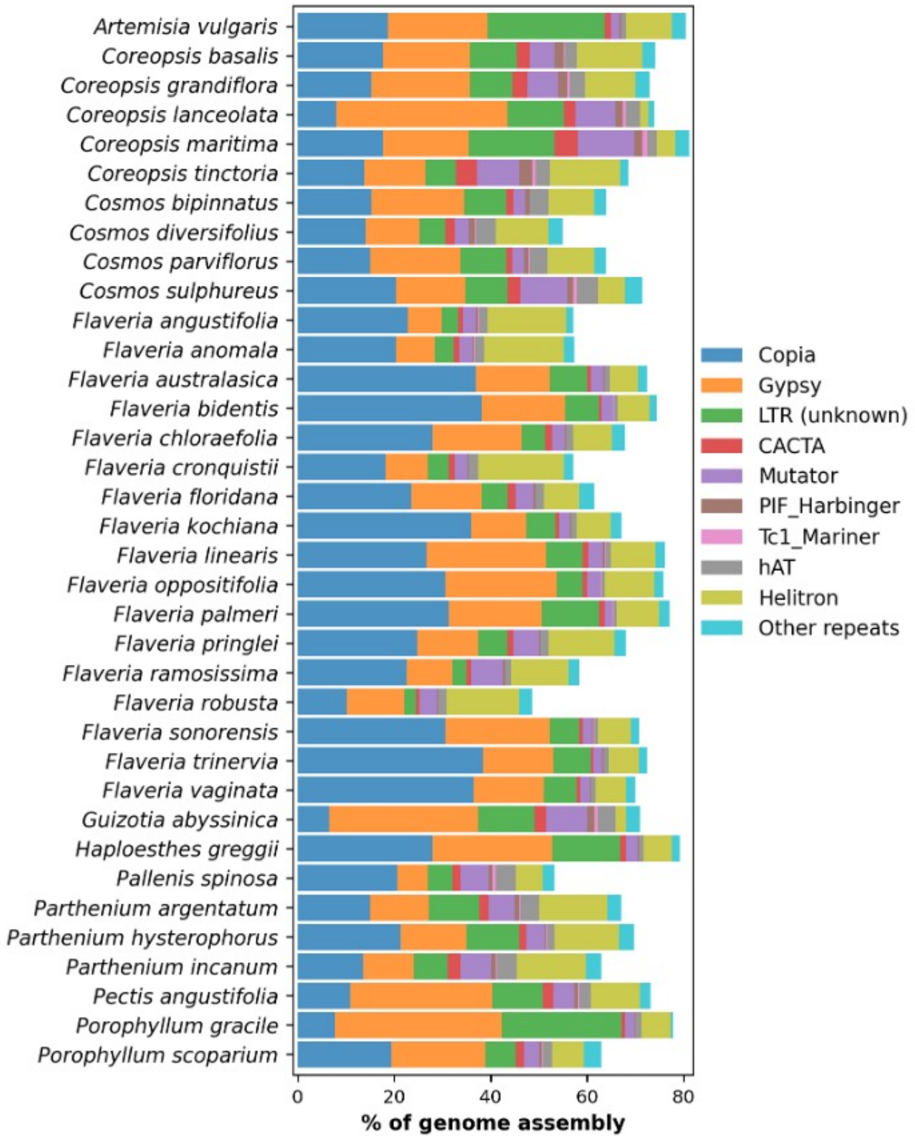
Artemisia vulgaris
Coreopsis basalis
Coreopsis grandiflora
Coreopsis lanceolata
Coreopsis maritima
Coreopsis tinctoria
Cosmos bipinnatus
Cosmos diversifolius
Cosmos parviflorus
Cosmos sulphureus
Flaveria angustifolia
Flaveria anomala
Flaveria australasica
Flaveria bidentis
Flaveria chloraefolia
Flaveria cronquistii
Flaveria floridana
Flaveria kochiana
Flaveria linearis
Flaveria oppositifolia
Flaveria palmeri
Flaveria pringlei
Flaveria ramosissima
Flaveria robusta
Flaveria sonorensis
Flaveria trinervia
Flaveria vaginata
Guizotia abyssinica
Haploesthes greggii
Pallenis spinosa
Parthenium argentatum
Parthenium hysterophorus
Parthenium incanum
Pectis angustifolia
Porophyllum gracile
Porophyllum scaparium
0
20
40
60
80
% of genome assembly
Copia
Gypsy
LTR (unknown)
CACTA
Mutator
PIF_Harbinger
Tc1_Mariner
hAT
Helitron
Other repeats
