## Supplementary Figures for "Ancestral polyploidy and non-linear dosage shifts define the photosynthetic transition in the Heliantheae alliance": Figure.S4.pdf

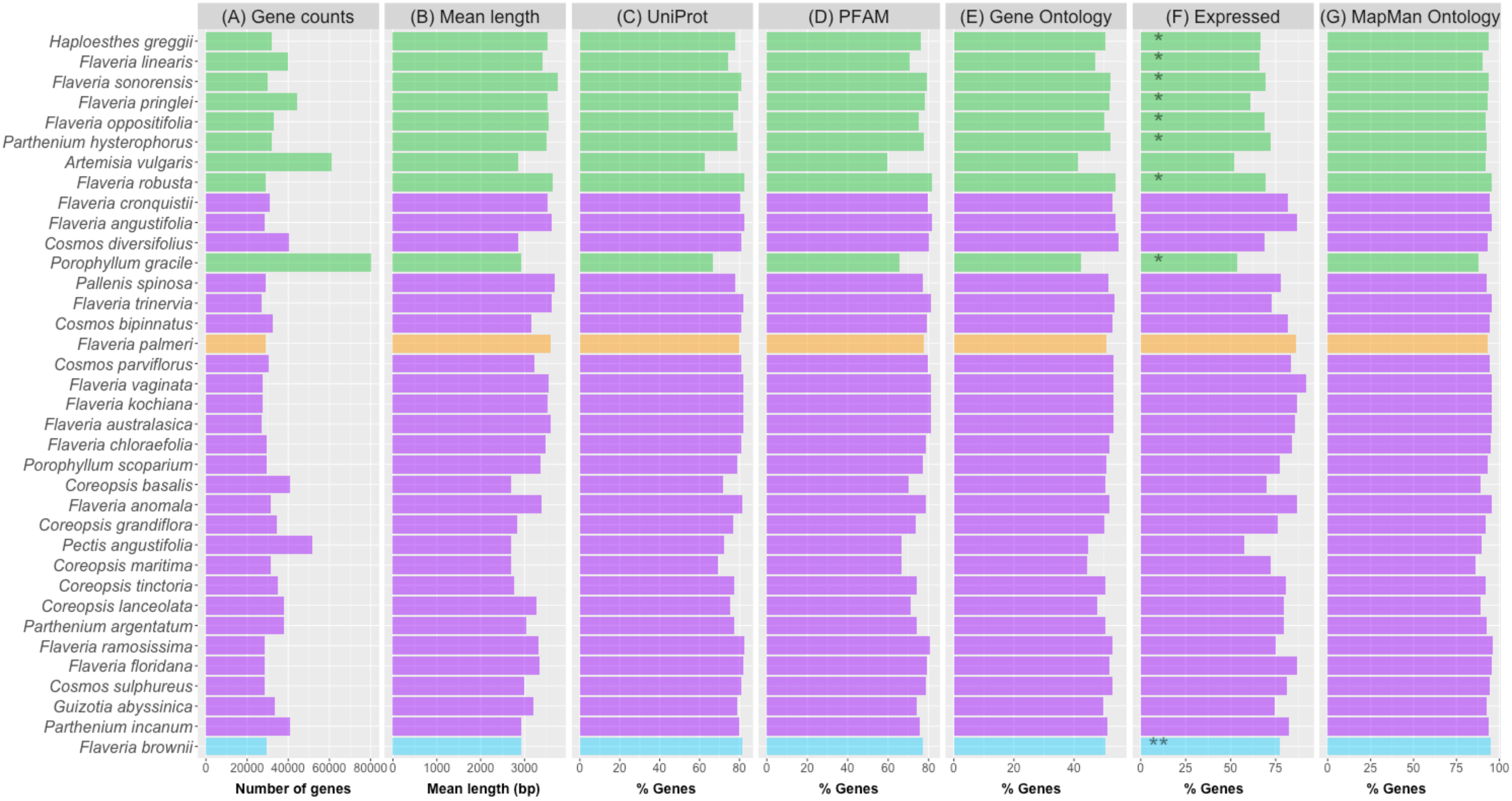

(A) Gene counts
(B) Mean length
(C) UniProt
(D) PFAM
(E) Gene Ontology
(F) Expressed
(G) MapMan Ontology
Haploesthes greggii
Flaveria linearis
Flaveria sonorensis
Flaveria pringlei
Flaveria oppositifolia
Parthenium hysterophorus
Artemisia vulgaris
Flaveria robusta
Flaveria cronquistii
Flaveria angustifolia
Cosmos diversifolius
Porophyllum gracile
Pallenis spinosa
Flaveria trinervia
Cosmos bipinnatus
Flaveria palmeri
Cosmos parviflorus
Flaveria vaginata
Flaveria kochiana
Flaveria australasica
Flaveria chloraefolia
Porophyllum scoparium
Coreopsis basalis
Flaveria anomala
Coreopsis grandiflora
Pectis angustifolia
Coreopsis maritima
Coreopsis tinctoria
Coreopsis lanceolata
Parthenium argentatum
Flaveria ramosissima
Flaveria floridana
Cosmos sulphureus
Guizotia abyssinica
Parthenium incanum
Flaveria brownii
0 20000 40000 60000 80000
Number of genes
0 1000 2000 3000
Mean length (bp)
0 20 40 60 80
% Genes
0 20 40 60 80
% Genes
0 20 40
% Genes
0 25 50 75
% Genes
0 25 50 75 100
% Genes
