## Supplementary Figures for "Ancestral polyploidy and non-linear dosage shifts define the photosynthetic transition in the Heliantheae alliance": Figure.S5.pdf

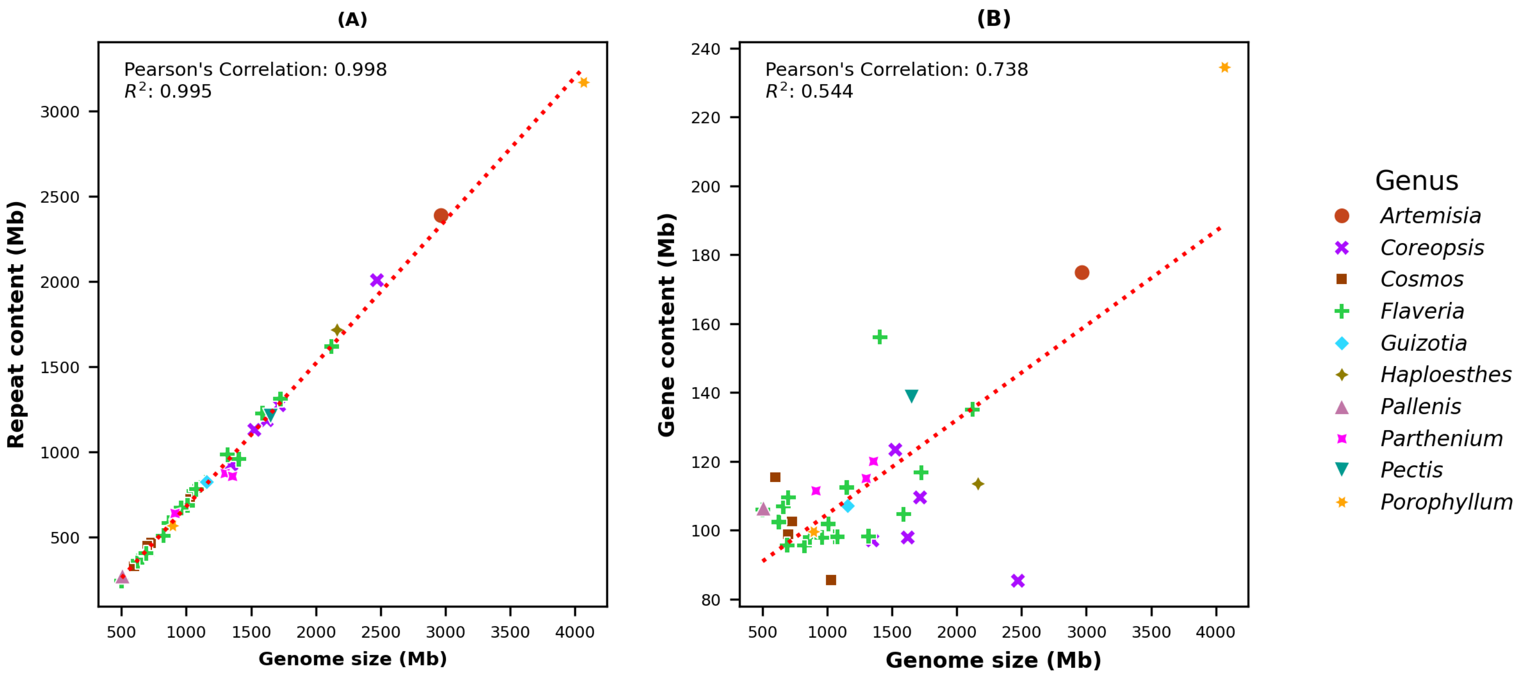

(A)
Pearson's Correlation: 0.998
$R^2$: 0.995
Repeat content (Mb)
Genome size (Mb)
(B)
Pearson's Correlation: 0.738
$R^2$: 0.544
Gene content (Mb)
Genome size (Mb)
Genus
Artemisia
Coreopsis
Cosmos
Flaveria
Guizotia
Haploesthes
Pallenis
Parthenium
Pectis
Porophyllum
