## Supplementary Figures for "Ancestral polyploidy and non-linear dosage shifts define the photosynthetic transition in the Heliantheae alliance": Figure.S6.pdf

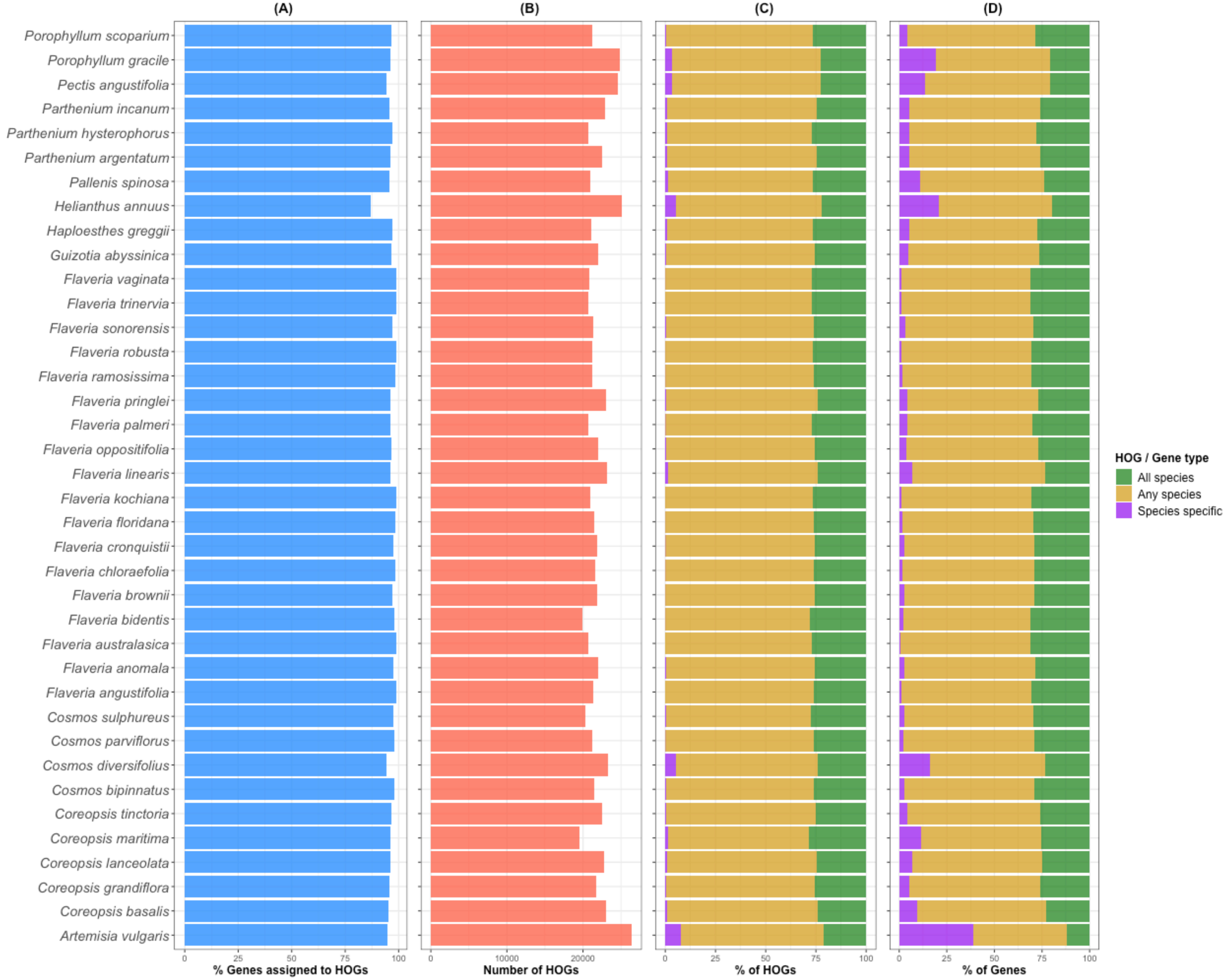
(A)
(B)
(C)
(D)
Porophyllum scoparium
Porophyllum gracile
Pectis angustifolia
Parthenium incanum
Parthenium hysterophorus
Parthenium argentatum
Pallenis spinosa
Helianthus annuus
Haploesthes greggii
Guizotia abyssinica
Flaveria vaginata
Flaveria trinervia
Flaveria sonorensis
Flaveria robusta
Flaveria ramosissima
Flaveria pringlei
Flaveria palmeri
Flaveria oppositifolia
Flaveria linearis
Flaveria kochiana
Flaveria floridana
Flaveria cronquistii
Flaveria chloraefolia
Flaveria brownii
Flaveria bidentis
Flaveria australasica
Flaveria anomala
Flaveria angustifolia
Cosmos sulphureus
Cosmos parviflorus
Cosmos diversifolius
Cosmos bipinnatus
Coreopsis tinctoria
Coreopsis maritima
Coreopsis lanceolata
Coreopsis grandiflora
Coreopsis basalis
Artemisia vulgaris
0 25 50 75 100
% Genes assigned to HOGs
0 10000 20000
Number of HOGs
0 25 50 75 100
% of HOGs
0 25 50 75 100
% of Genes
HOG / Gene type
All species
Any species
Species specific
