## Supplementary Figures for "Ancestral polyploidy and non-linear dosage shifts define the photosynthetic transition in the Heliantheae alliance": Figure.S7.pdf

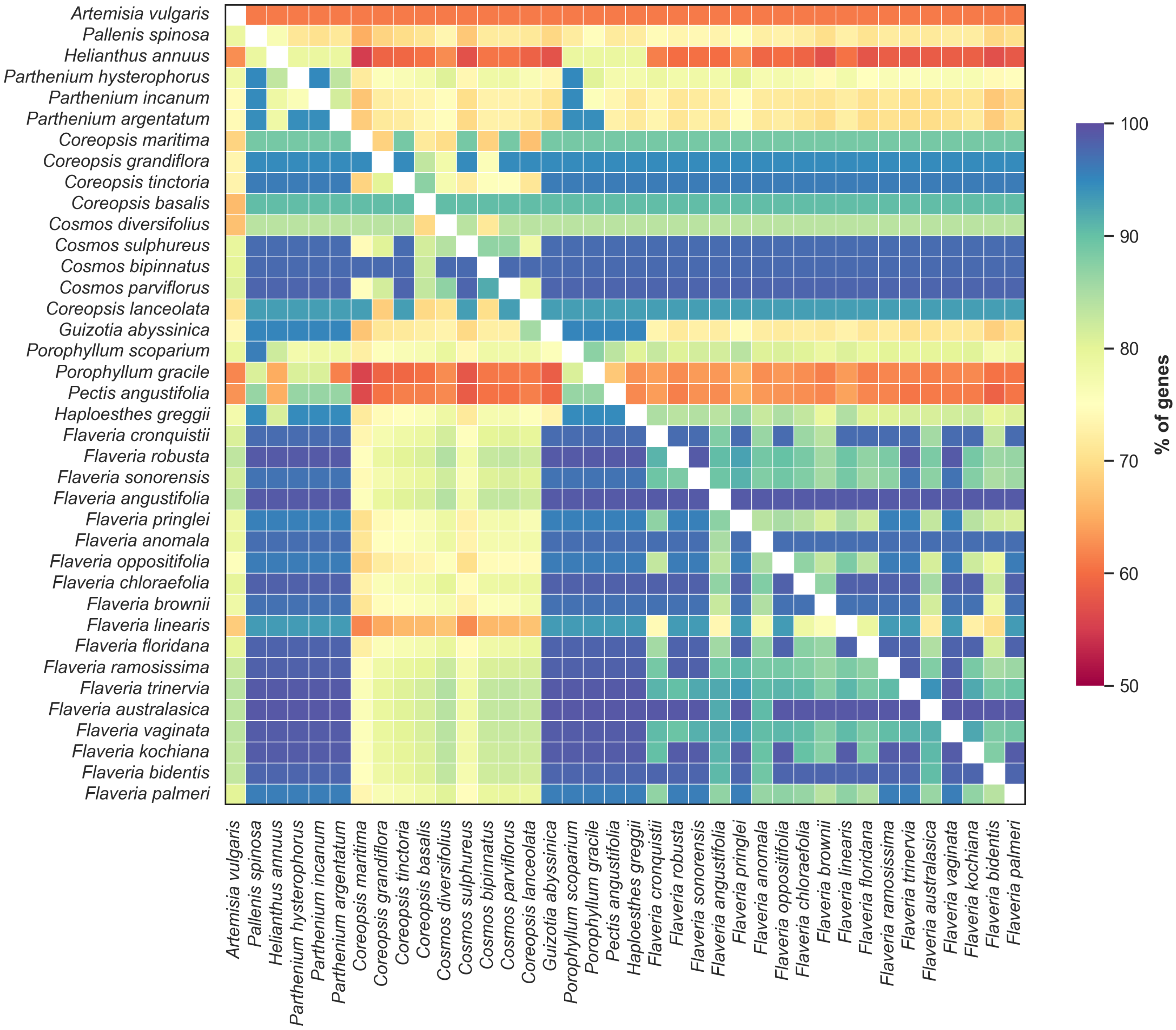

Artemisia vulgaris
Pallenis spinosa
Helianthus annuus
Parthenium hysterophorus
Parthenium incanum
Parthenium argentatum
Coreopsis maritima
Coreopsis grandiflora
Coreopsis tinctoria
Coreopsis basalis
Cosmos diversifolius
Cosmos sulphureus
Cosmos bipinnatus
Cosmos parviflorus
Coreopsis lanceolata
Guizotia abyssinica
Porophyllum scoparium
Porophyllum gracile
Pectis angustifolia
Haploesthes greggii
Flaveria cronquistii
Flaveria robusta
Flaveria sonorensis
Flaveria angustifolia
Flaveria pringlei
Flaveria anomala
Flaveria oppositifolia
Flaveria chloraefolia
Flaveria brownii
Flaveria linearis
Flaveria floridana
Flaveria ramosissima
Flaveria trinervia
Flaveria australasica
Flaveria vaginata
Flaveria kochiana
Flaveria bidentis
Flaveria palmeri
100
90
80
70
60
50
% of genes
