## Supplementary Figures for "Ancestral polyploidy and non-linear dosage shifts define the photosynthetic transition in the Heliantheae alliance": Figure.S8.pdf

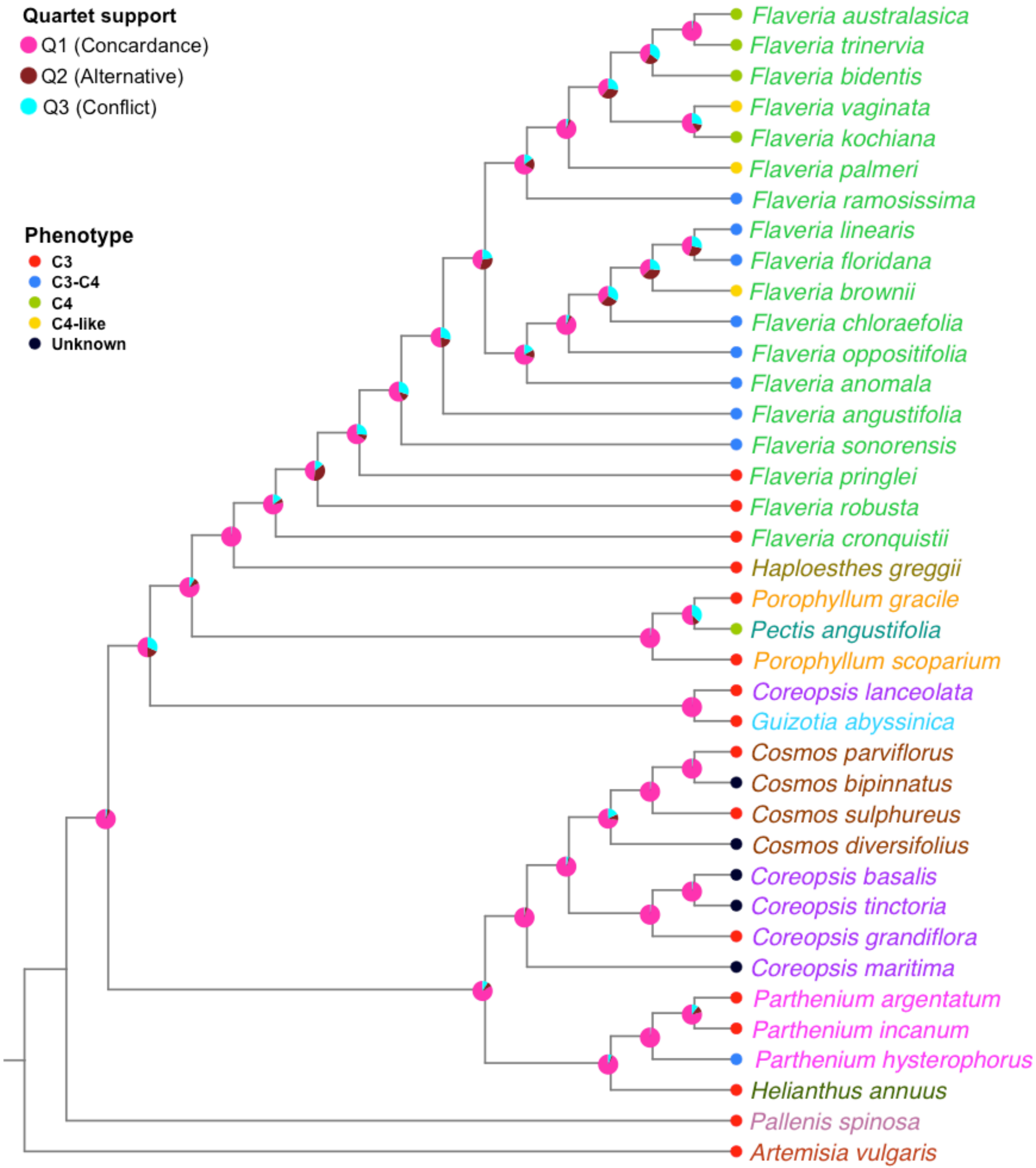
Quartet support
Q1 (Concardance)
Q2 (Alternative)
Q3 (Conflict)
Phenotype
C3
C3-C4
C4
C4-like
Unknown
Flaveria australasica
Flaveria trinervia
Flaveria bidentis
Flaveria vaginata
Flaveria kochiana
Flaveria palmeri
Flaveria ramosissima
Flaveria linearis
Flaveria floridana
Flaveria brownii
Flaveria chloraefolia
Flaveria oppositifolia
Flaveria anomala
Flaveria angustifolia
Flaveria sonorensis
Flaveria pringlei
Flaveria robusta
Flaveria cronquistii
Haploesthes greggii
Porophyllum gracile
Pectis angustifolia
Porophyllum scoparium
Coreopsis lanceolata
Guizotia abyssinica
Cosmos parviflorus
Cosmos bipinnatus
Cosmos sulphureus
Cosmos diversifolius
Coreopsis basalis
Coreopsis tinctoria
Coreopsis grandiflora
Coreopsis maritima
Parthenium argentatum
Parthenium incanum
Parthenium hysterophorus
Helianthus annuus
Pallenis spinosa
Artemisia vulgaris
