## Supplementary Figures for "Ancestral polyploidy and non-linear dosage shifts define the photosynthetic transition in the Heliantheae alliance": Figure.S11.pdf

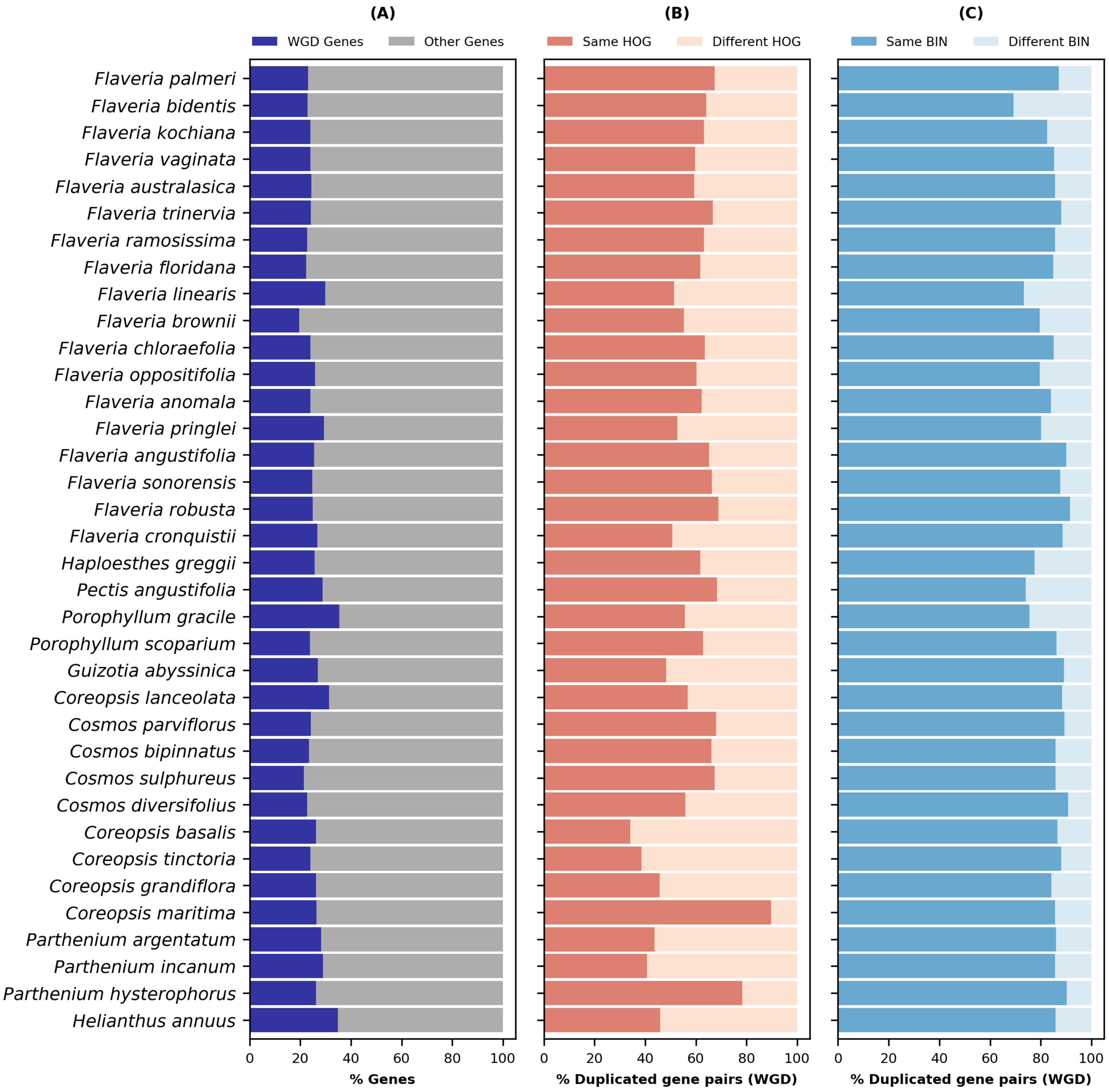
(A)
WGD Genes
Other Genes
(B)
Same HOG
Different HOG
(C)
Same BIN
Different BIN
Flaveria palmeri
Flaveria bidentis
Flaveria kochiana
Flaveria vaginata
Flaveria australasica
Flaveria trinervia
Flaveria ramosissima
Flaveria floridana
Flaveria linearis
Flaveria brownii
Flaveria chloraefolia
Flaveria oppositifolia
Flaveria anomala
Flaveria pringlei
Flaveria angustifolia
Flaveria sonorensis
Flaveria robusta
Flaveria cronquistii
Haploesthes greggii
Pectis angustifolia
Porophyllum gracile
Porophyllum scoparium
Guizotia abyssinica
Coreopsis lanceolata
Cosmos parviflorus
Cosmos bipinnatus
Cosmos sulphureus
Cosmos diversifolius
Coreopsis basalis
Coreopsis tinctoria
Coreopsis grandiflora
Coreopsis maritima
Parthenium argentatum
Parthenium incanum
Parthenium hysterophorus
Helianthus annuus
0
20
40
60
80
100
% Genes
% Duplicated gene pairs (WGD)
% Duplicated gene pairs (WGD)
