## Supplementary Figures for "Ancestral polyploidy and non-linear dosage shifts define the photosynthetic transition in the Heliantheae alliance": Figure.S12.pdf

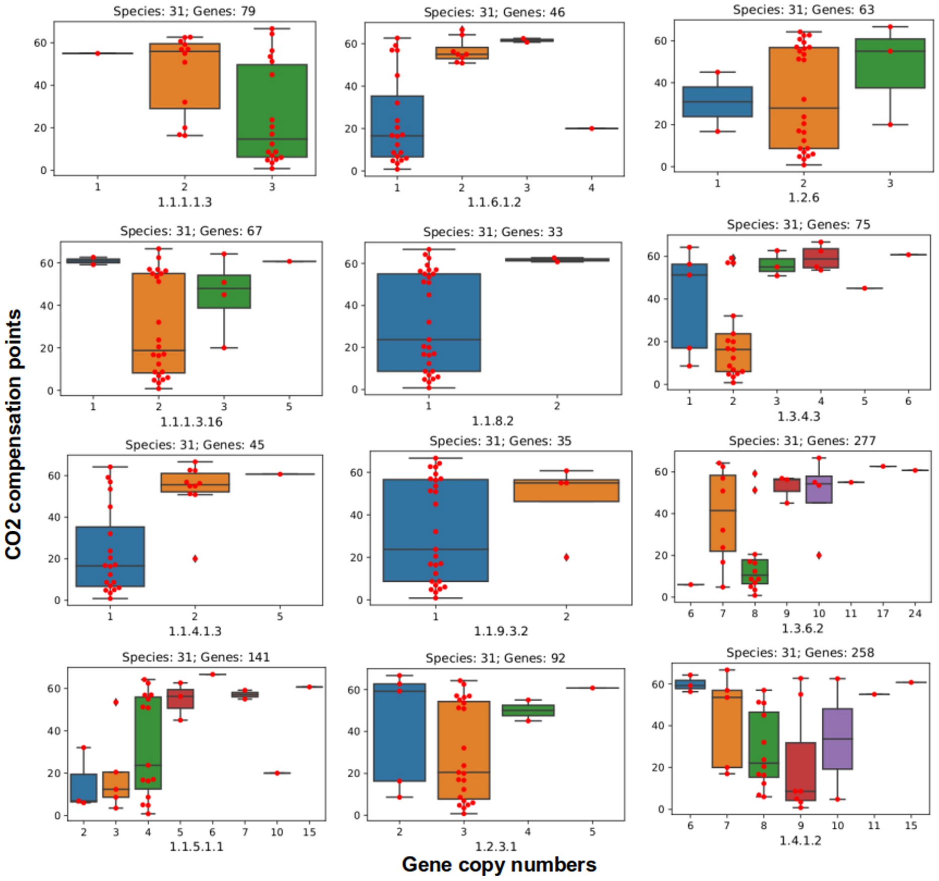
CO2 compensation points
Species: 31; Genes: 79
1.1.1.1.3
Species: 31; Genes: 46
1.1.6.1.2
Species: 31; Genes: 63
1.2.6
Species: 31; Genes: 67
1.1.1.3.16
Species: 31; Genes: 33
1.1.8.2
Species: 31; Genes: 75
1.3.4.3
Species: 31; Genes: 45
1.1.4.1.3
Species: 31; Genes: 35
1.1.9.3.2
Species: 31; Genes: 277
1.3.6.2
Species: 31; Genes: 141
1.1.5.1.1
Species: 31; Genes: 92
1.2.3.1
Species: 31; Genes: 258
1.4.1.2
Gene copy numbers
