## Supplementary Figures for "Ancestral polyploidy and non-linear dosage shifts define the photosynthetic transition in the Heliantheae alliance": Figure.S13.pdf

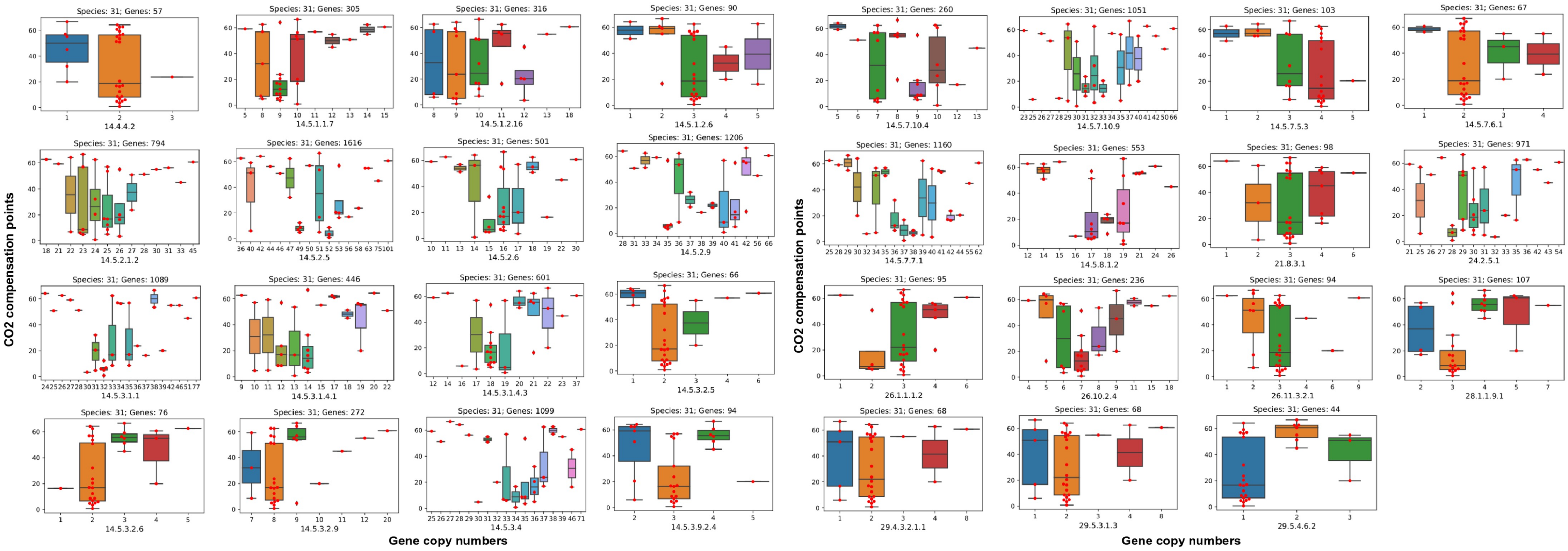

CO2 compensation points
Gene copy numbers
Species: 31; Genes: 57
14.4.4.2
Species: 31; Genes: 305
14.5.1.1.7
Species: 31; Genes: 316
14.5.1.2.16
Species: 31; Genes: 90
14.5.1.2.6
Species: 31; Genes: 794
14.5.2.1.2
Species: 31; Genes: 1616
14.5.2.5
Species: 31; Genes: 501
14.5.2.6
Species: 31; Genes: 1206
14.5.2.9
Species: 31; Genes: 1089
14.5.3.1.1
Species: 31; Genes: 446
14.5.3.1.4.1
Species: 31; Genes: 601
14.5.3.1.4.3
Species: 31; Genes: 66
14.5.3.2.5
Species: 31; Genes: 76
14.5.3.2.6
Species: 31; Genes: 272
14.5.3.2.9
Species: 31; Genes: 1099
14.5.3.4
Species: 31; Genes: 94
14.5.3.9.2.4
CO2 compensation points
Gene copy numbers
Species: 31; Genes: 260
14.5.7.10.4
Species: 31; Genes: 1051
14.5.7.10.9
Species: 31; Genes: 103
14.5.7.5.3
Species: 31; Genes: 67
14.5.7.6.1
Species: 31; Genes: 1160
14.5.7.7.1
Species: 31; Genes: 553
14.5.8.1.2
Species: 31; Genes: 98
21.8.3.1
Species: 31; Genes: 971
24.2.5.1
Species: 31; Genes: 95
26.1.1.1.2
Species: 31; Genes: 236
26.10.2.4
Species: 31; Genes: 94
26.11.3.2.1
Species: 31; Genes: 107
28.1.1.9.1
Species: 31; Genes: 68
29.4.3.2.1.1
Species: 31; Genes: 68
29.5.3.1.3
Species: 31; Genes: 44
29.5.4.6.2
