## Supplementary figures and images for "Ancestral polyploidy and non-linear dosage shifts define the photosynthetic transition in the Heliantheae alliance"

### Figure.S10.pdf

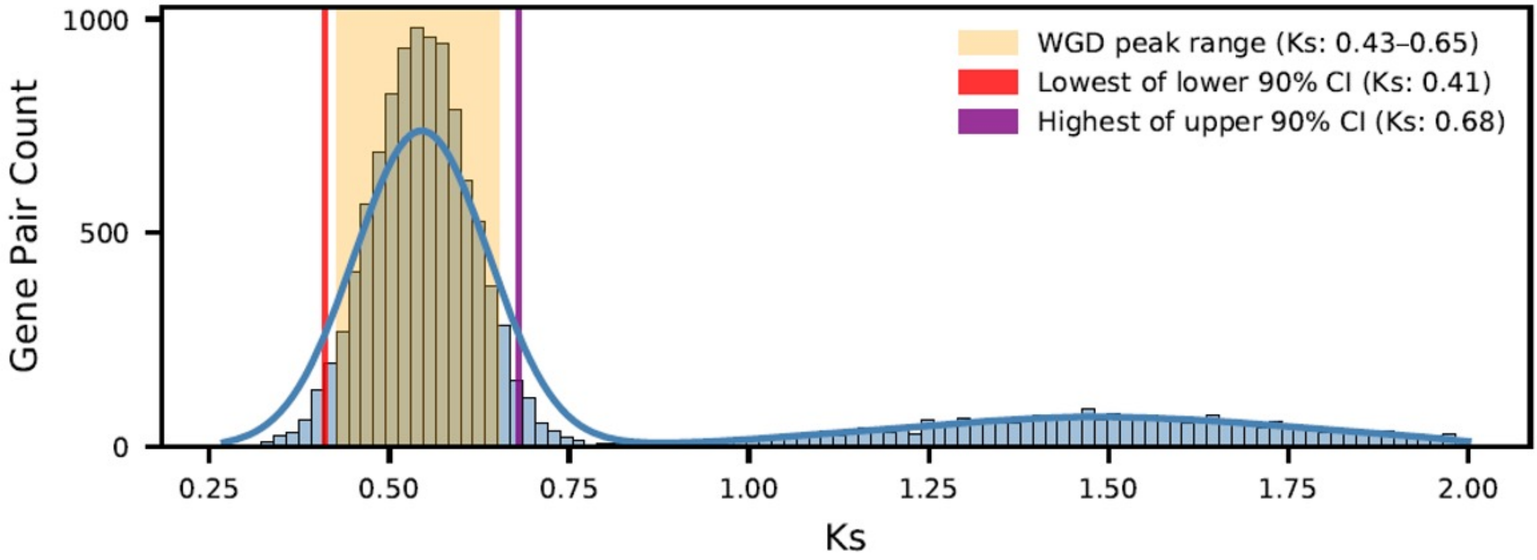

WGD peak range (Ks: 0.43–0.65)
Lowest of lower 90% CI (Ks: 0.41)
Highest of upper 90% CI (Ks: 0.68)
Gene Pair Count
Ks
0
500
1000
0.25
0.50
0.75
1.00
1.25
1.50
1.75
2.00
